# High resolution mapping of the actin fusion focus reveals myosin V-dependent transport of formin for actin aster compaction

**DOI:** 10.1101/2025.04.29.651207

**Authors:** Valentine Thomas, Hikari Mase, Laetitia Michon, Andrea Picco, Marko Kaksonen, Sophie G Martin

## Abstract

Many cellular processes such as polarized growth and secretion require the formation of specific actin networks. In fungi, the morphogenetic process of cell-cell fusion requires cell wall digestion mediated by the local secretion of lytic enzymes at the site of cell-cell contact. In *Schizosaccharomyces pombe*, lytic enzyme-containing secretory vesicles are transported by the myosin V Myo52 on the actin fusion focus, an aster-like actin network assembled by the condensate-forming formin Fus1. The fusion focus also concentrates proteins regulating cell polarity, cell-cell communication, actin cytoskeleton, exocytosis and membrane merging, but their precise position from the time of focus formation to cell fusion is unknown. Here, using centroid tracking and averaging, we present a first spatiotemporal map of the fusion site with a precision of 8 nm. We show that the bulk of secretory vesicles remains at constant distance from the plasma membrane as the actin structure condenses. Notably, though necessary to transport vesicles, Myo52 detaches from the vesicle pool and colocalizes with Fus1 in a more membrane-proximal position. We show that Myo52 physically interacts with Fus1 and transports it along actin filaments, and that Myo52 and Fus1 actin assembly activity contribute to focus compaction. Thus, myosin V-driven transport of the formin Fus1 along actin filaments nucleated by other Fus1 molecules underlies a positive feedback mechanism for actin aster formation.

## Introduction

Cell-cell fusion is a ubiquitous process, which underlies fertilization for sexual reproduction and the formation of various tissues during development. In fungi, the cell wall represents a major barrier to cell fusion and must be locally digested to allow membrane merging. In yeast cells, where the process has been best studied, local cell wall degradation requires the formin-dependent assembly at the site of cell-cell contact of a linear actin filament structure, named the actin fusion focus ^1^. The actin fusion focus serves to concentrate type V myosin motors, which transport secretory vesicles containing cell wall lytic enzymes. Local content release leads to local piercing of the cell wall. Thus, the early steps of cell-cell fusion in yeast represent a fascinating case of hyper-polarized secretion.

The core mechanisms of polarized secretion are conserved across organisms and play pivotal roles in various biological systems, from the activity of neuronal synapses to the polarized growth of walled cells. Detailed mechanistic understanding largely comes from work on polarized mitotic growth in yeasts (mostly *S. cerevisiae*, but also *S. pombe*). Secretory vesicles, marked by the Rab GTPases Ypt3 (Rab11-like) and Sec4 (Rab8-like) as well as by Sec4’s guanine nucleotide exchange factor activator Sec2, first form at the trans-Golgi network ^2^. Immediately upon formation, they are bound by myosin V through direct interactions of its C-terminal cargo-binding domain to the Rab GTPases ^3–7^. Myosin V-driven movement then transports them to the site of polarized growth along actin cables nucleated by formin proteins ^8–10^. The octameric exocyst complex is recruited to the vesicle during transport ^2, 11^, likely through direct binding of its Sec15 subunit with the myosin V ^6^, and then tethers the incoming vesicle at the plasma membrane. In the 5 seconds between initial tethering and vesicle fusion, the Sec4 GEF departs, followed by the myosin, before the SNARE complex mediates membrane fusion ^2, 12, 13^.

The actin cytoskeleton plays a critical role in defining the site of polarized secretion. In the complex cellular F-actin landscape, myosin V selectively moves on linear actin filaments decorated by tropomyosin ^14, 15^. In large cells, such as mouse oocytes, nucleators of linear actin filaments (Spir; formin-2) assemble a network of filaments from both vesicles and the plasma membrane, allowing their myosin V-dependent collective movements towards the cell surface ^16^. In small yeast cells, formins assemble actin cables from the poles of the cell (or bud site), which traverse the cell and are used as long-range linear tracks by myosin V ^17^. Forces exerted by myosin V during transport also contribute to actin cable extension ^18^.

During sexual reproduction in fission yeast *Schizosaccharomyces pombe*, assembly of the actin fusion focus by the formin Fus1, expressed solely during mating, is strictly required for cell fusion ^19–21^. The fusion focus, an aster-like organization of linear actin filaments, serves to concentrate 60-nm wide vesicles containing cell wall lytic enzymes, transported by the ubiquitous myosin V Myo52 ^9, 19, 22, 23^. Focus assembly occurs at the interface of P and M cell pairs, and its stabilization at a position facing the partner is tightly linked to the pheromone-Ras-MAPK signaling that takes place between partner cells. Indeed, enrichment of pheromone-Ras-MAPK signaling on the fusion focus leads to its stabilization, which is prevented at early time points by global inactivation of Ras ^24, 25^. This coupling ensures that local cell wall digestion occurs at the middle of the contact zone, preserving cell integrity.

The set-up is largely symmetric between mating partners: each of the two cell types assembles a focus at the cell-cell contact site, and a single focus can, though inefficiently, promote cell wall erosion ^19^. However, some asymmetries have been observed, where the M-cell stabilizes the structure first and is more protrusive than the P-cell ^19, 23^.

Several mechanisms contribute to focusing the fusion focus. Most prominently, Fus1 bears an intrinsically disordered domain (IDR), thought to mediate weak multivalent self-interactions that underlie the formation of a biomolecular condensate. This was shown through the functional replacement of this domain with heterologous domains exhibiting liquid-liquid phase separation in vitro ^26^. Deletion of Fus1 IDR specifically alters the ability of Fus1 to condense in a focus, and Fus1^ΔIDR^ decorates a significantly wider area at the plasma membrane and does not support cell-cell fusion. A second mechanism involves a single-headed type V myosin, named Myo51, with no role in vesicular transport ^27^. Myo51 recruits the coiled-coil proteins Rng8 and Rng9 to the fusion focus, where these proteins promote its coalescence likely through cross-linking of tropomyosin-decorated filaments ^28^. The dimeric myosin V Myo52 responsible for vesicular transport may also play an unknown role because Fus1 was shown to occupy a wider area at the plasma membrane when both Myo51 and Myo52 were deleted ^19^.

The fusion focus not only promotes cell wall piercing, which is the kinetically slow step during cell fusion ^23^, but also concentrates many proteins regulating cell polarity, pheromone signaling, actin cytoskeleton organization, exocytosis, or transmembrane proteins required for the later step of plasma membrane merging, including the conserved fungal protein Prm1 ^19, 24, 25, 28–35^. Whether these proteins occupy distinct positions at the fusion site, or whether they are mixed in the fusion focus, as perhaps suggested by its condensate properties, is unknown. Because the dimensions of the fusion site are of only a few 100 nm, as observed by electron tomography ^23^, and the fusion focus matures over time ^19^, a live-cell super-resolution approach is required to resolve the position of individual proteins.

Here, by developing a method combining fluorescence centroid tracking with spatial and temporal references, we establish a high-precision median map of fusion focus components over time. We find that individual proteins exhibit distinct trajectories, indicating stereotypical order in the fusion focus. The map provides new insights in polarized secretion and actin organization. Notably, we find that secretory vesicles form a pool that remains at constant distance from the plasma membrane. While Myo52 is responsible for their transport, the myosin detaches from them and colocalizes with Fus1 at a more membrane-proximal position during focus maturation. We show interaction of Myo52 and Fus1 through co-immunoprecipitation and demonstrate that the formin is a Myo52 cargo. Consistent with Myo52 transporting Fus1, both Myo52 and Fus1 activity are required to restrict the lateral distribution of the formin. We propose that the association of Fus1 with Myo52 underlies positive feedback for Fus1 convergence. The myosin V Myo52 transports the formin Fus1 along filaments nucleated by other Fus1 dimers, leading to their focusing and formation of an actin aster.

## Results

### High-precision centroid median mapping method by live-cell epifluorescence imaging

To investigate the stereotypical organization of the fusion focus during mating, we aimed to obtain a high-precision spatiotemporal map, where the trajectory of each focus component is positioned over time relative to a constant reference. We adapted a method previously developed to map endocytic proteins in budding yeast cells ^36^, extending it to three-color live-cell imaging. Briefly, the method relies on precise detection of the centroid of fluorescence of any protein of interest, whose spatial and temporal coordinates are recorded relative to spatial and temporal references. High precision derives from averaging hundreds of traces, leading to a trajectory that describes the stereotypical behavior of the center of mass of the protein pool during cell fusion.

Because the type V myosin Myo52 is present throughout cell fusion and exhibits a strong fluorescence signal, we chose this protein as spatial reference, tagged with mScarlet-I in the cell where the map was measured, and with mTagBFP2 in the partner cell (Figure 1A). An axis passing through both Myo52 signals defines the fusion axis. We initially chose tdTomato to tag Myo52, but observed significant photoconversion into the green channel, motivating our switch to the more photostable mScarlet-I (Figure S1A-B). The partner cell further expressed a stable weakly fluorescent cytosolic protein (mRaspberry; ^37^), which was used as temporal reference to define fusion time, as it rapidly diffuses into the ‘map’ cell upon fusion pore opening. Monitoring the cytoplasmic signal in the ‘map’ cell revealed a steep increase in fluorescence, chosen as fusion time t = 0 (Figure 1B). Proteins of interest were tagged with sfGFP (or other green-fluorescent derivatives) and expressed in the same cell as Myo52-mScarlet-I, allowing to position them along the fusion axis (Table S1). This setup allowed three-color acquisition with a standard epifluorescence microscope.

**Figure 1:**
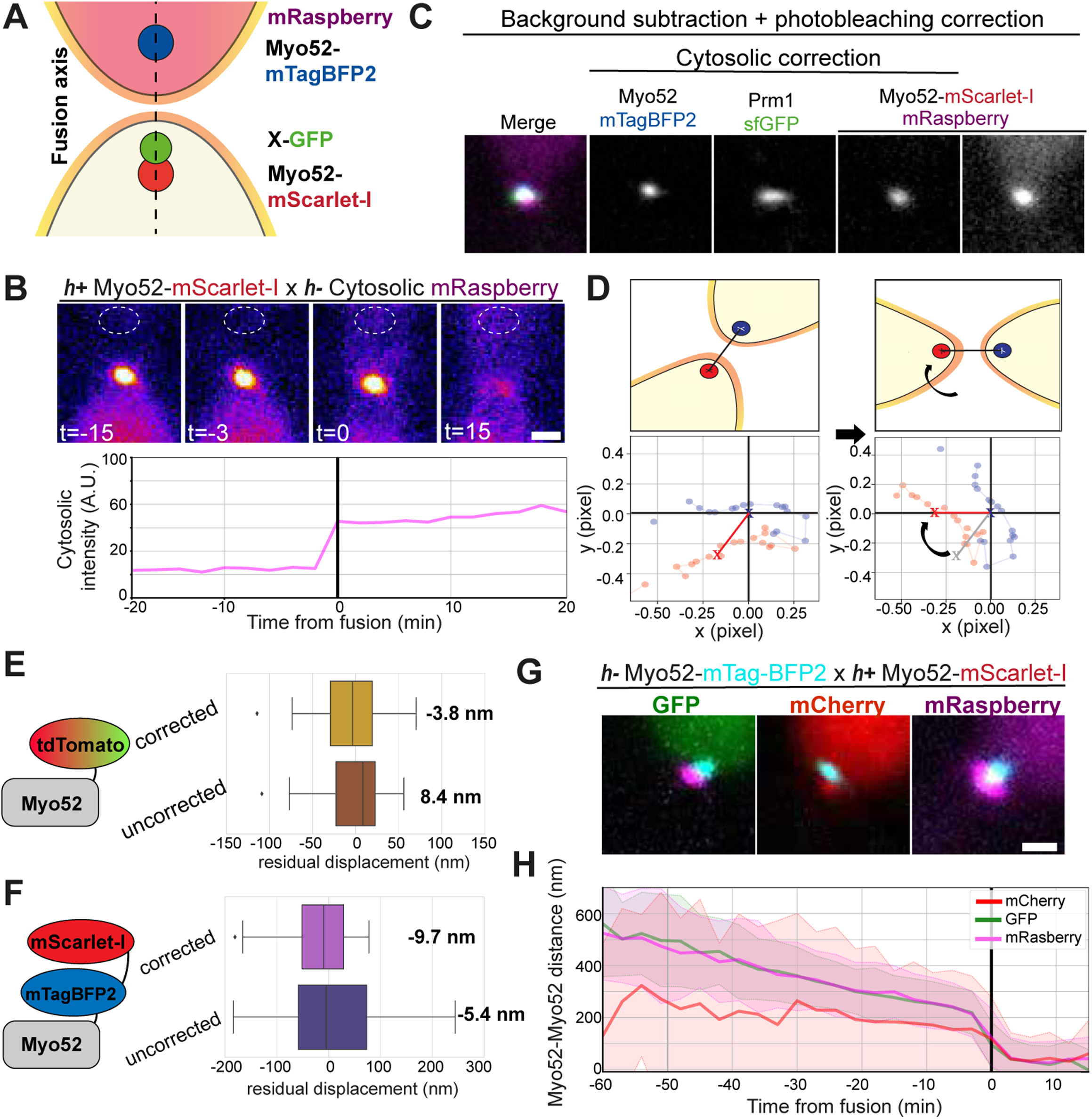
Imaging pipeline for high-resolution spatiotemporal mapping of fluorescence in the fusion focus. **A.** Scheme of the experimental system used for fusion focus mapping. **B.** Timelapse of RFP channel at the time of cell-cell fusion, showing cytoplasmic mRaspberry and Myo52-mScarlet-I signals. The dotted circle indicates the site of cytosolic fluorescence measurements displayed in the bottom graph. The black line indicates fusion time t = 0. **C.** Representative post-processing images of fusion focus components used for centroid tracking from BFP, GFP and RFP channels. All images were corrected for background and photobleaching. The middle three were also corrected for cytosolic noise. **D.** Illustrative scheme of trajectories spatial alignment. The fusion axis is defined between both Myo52’s trajectories center of mass, with that of *h-* cell defined as origin. It is then rotated around the origin to the x-axis. **E-F.** Schematics of constructs used to assess residual error between red and green (E; photoconverting tdTomato), and red and blue signals (F). The graphs show the residual displacement between fluorophores measured along the fusion axis before and after chromatic aberration correction. Box plots show median value and interquartile range, and whiskers maximum and minimum. **G.** Merge images showing *h+* cells expressing Myo52-mScarlet-I and *h-* cells co-expressing Myo52-mTagBFP2 and the indicated cytoplasmic fluorescent protein. **H.** The graph shows the median directed distance between Myo52 foci during the fusion process in cells as in (G). Median trajectories measured from cells with GFP or mRaspberry cytosolic marker are not significantly different (p-value > 0.05). Shaded zones show standard deviation. Black line shows fusion time. Scale bars are 1 µm.

Three-color imaging of the mating process was performed in duplicate for each strain and optimized to limit photobleaching while maximizing the number of fusion events recorded (see methods). Each channel was processed for signal isolation before centroid tracking as previously described ^36^. Briefly, the background signal was removed and photobleaching was corrected, before any remaining cytosolic signal was subtracted, thus isolating the focus signal (Figure 1C), whose centroid was tracked (see methods).

To spatially align trajectories, we used the center of mass of the Myo52 trajectory in the M cell, where the focus stabilizes first ^19^, as the origin for a coordinate system. All trajectories were then rotated around the origin such that the fusion axis – the line passing through both Myo52 trajectories’ center of mass – is on the x-axis (Figure 1D). The distances between Myo52-mScarlet-I and the GFP-tagged proteins of interest were measured for each time point along this axis, temporally aligned to t0, and used to obtain median protein trajectories throughout the fusion process.

There are two major sources of error in centroid measurements. First, even though yeast cells are restricted to a quasi-two-dimensional plane by our imaging strategy, displacement in z of the fusion focus would lead to measurement of a projected distance in x-y, yielding a value below reality. We reduced this source of error by manually curating trajectories, to remove noise and ensure that trajectories for all proteins were present. For large distances between Myo52 foci at the start of the fusion process, we estimate that imprecision in z yields a roughly 2% error in distance (i.e. ∼1 nm per 50 nm measured distance; see methods). Second, chromatic aberrations are a source of error when comparing fluorophores with distinct emission wavelengths. We thus corrected trajectory coordinates for lateral chromatic aberration (see methods). We then assessed the accuracy of the method by applying it to Myo52 tagged with two colors and measuring the distance between the two signals. We took advantage of tdTomato red-to-green photoconversion to measure any green-red shift. We constructed a double-tagged Myo52-mtagBFP2-mScarlet-I (with Myo52-GFP in the partner cell) to assess any blue-red shift. Chromatic aberration correction reduced the residual error to an average of 4 nm between green and red, and decreased measurement variability, with an average of 10 nm between blue and red channels (Figure 1E-F). A conservative estimate of the total residual error in distances between red and green signals is ∼ 8 nm (see methods).

Given our 3-channels imaging strategy, we did not have an independent channel to image the cytosolic marker used for temporal alignment. We thus verified its presence did not lead to a lateral displacement of centroid position after cytosolic signal subtraction. We first measured the median distance between Myo52 signals without interference from cytosolic signal, using GFP as cytosolic marker. As in previous manual measurements ^19^, the distance between fusion foci progressively shortened, until abrupt convergence at fusion time (Figure 1G-H). We then evaluated the red-fluorescent cytosolic proteins mCherry and mRaspberry. Use of mCherry, which exhibits high fluorescence intensity relative to Myo52-mScarlet-I, led to significant reduction in the Myo52-Myo52 distance, indicating displacement of the Myo52-mScarlet-I centroid towards the partner cell. The measure was also more variable indicated by a large standard deviation. By contrast, cytosolic mRaspberry, which exhibits weaker fluorescence, did not affect the median distance between Myo52 foci, which was indistinguishable from the reference obtained with cytosolic GFP (Figure 1G-H). We conclude that subtraction of the cytosolic mRaspberry signal does not alter the Myo52-mScarlet-I signal. We thus used mRaspberry as cytosolic marker in all further measurements.

The centroid-tracking method described above can be applied to signals that are (roughly) spherical. However, some of the GFP-tagged proteins of interest exhibit a more extended distribution at the cell-cell interface. In these cases, instead of measuring the centroid of the GFP signal, we measured the GFP fluorescence profile along the fusion axis between the Myo52 centroids and fitted it with a Gaussian function to define the center of mass of the GFP signal along the fusion axis (see methods). When both methods were directly compared on the same imaging dataset, they yielded overlapping trajectories (Figure S1C).

Thus, we established a method to measure the median position of any protein at the fusion site along the fusion axis with an error ∼ 8 nm.

### Stereotypical architecture and maturation of the fusion focus

We initially compared the trajectories of five proteins with roles in cell polarization, cytoskeletal organization, exocytosis and membrane fusion from 60 min before until 15 min after fusion. We represent their median oriented distance to the Myo52-mScarlet-I reference in the same *h+* cell (Figure 2A). This analysis revealed that each of these proteins exhibited a trajectory distinct from the others. Scd2, a scaffold protein associated with active Cdc42 GTPase at the plasma membrane ^38, 39^, and Prm1, a transmembrane protein involved in membrane merging ^29^, localized proximal to the partner cell relative to the Myo52 reference. Myo51, a myosin that decorates actin filaments and plays structural cross-linking roles ^28^, localized distally in the cytosol and moved progressively closer to the Myo52 reference. The formin Fus1 and exocyst subunit Exo84 localized in an intermediate zone, but their trajectories showed distinct slopes. Thus, individual proteins in the fusion focus exhibit distinct trajectories.

**Figure 2:**
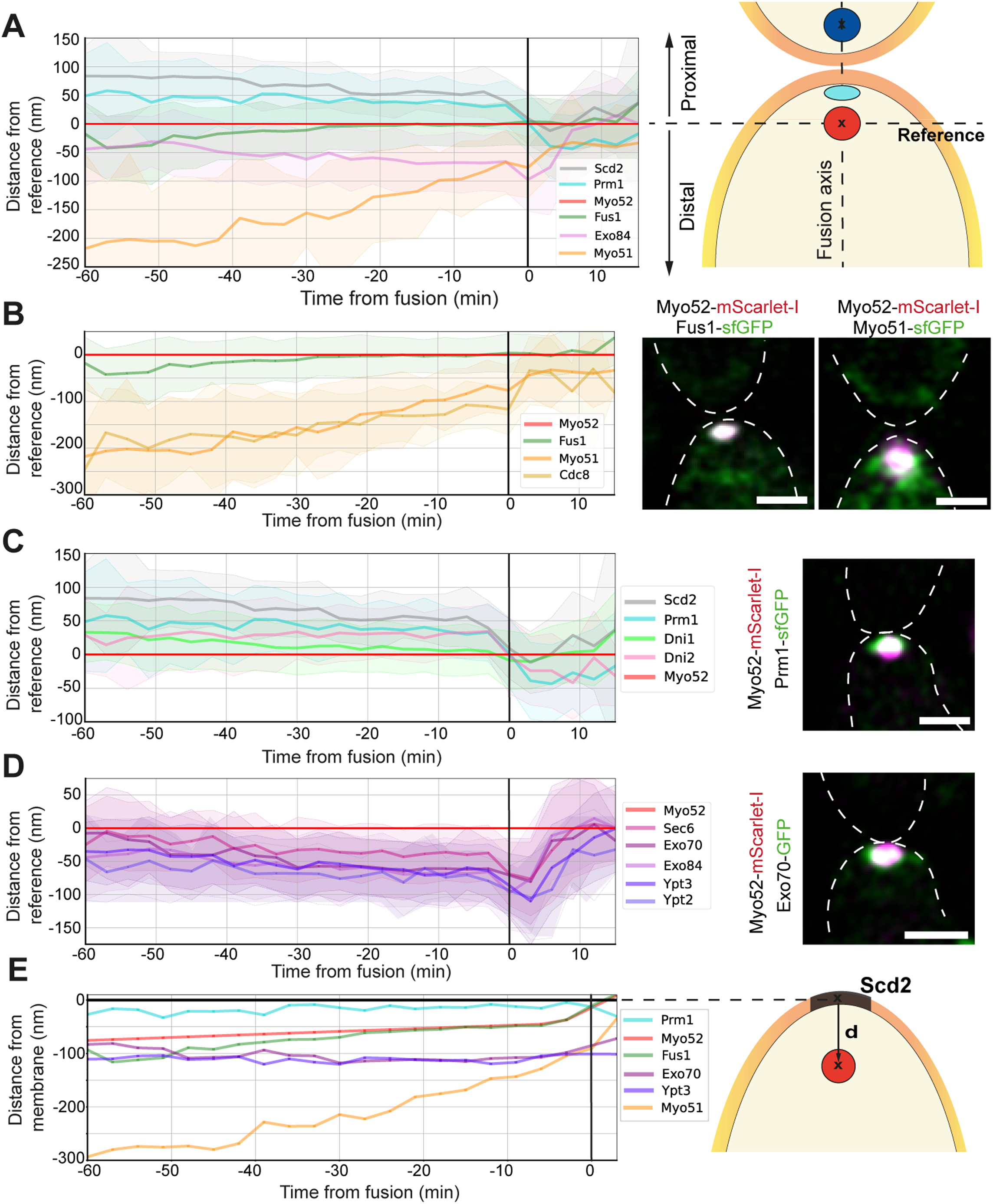
Spatiotemporal map of the fusion focus. **A.** Median directed distances of Scd2, Prm1, Fus1, Exo84 and Myo51 from Myo52 over the fusion process. The scheme on the right provides orientation, with positive values indicating position proximal to the partner cell and negative values position more cytosolic than Myo52. **B-D.** Median directed distances from Myo52 of (B) actin regulators, (C) plasma membrane-associated factors and (D) exocyst and Rab GTPases labelling secretory vesicles. Airyscan images of selected proteins are shown on the right. **E.** Transposed directed distances from Scd2, representing the plasma membrane, with explanatory scheme on the right. On all graphs, shaded zones show standard deviation, black line shows fusion time. Scale bars are 1 µm.

As asymmetries between mating types have been described in terms of fusion focus focalization timing as well as membrane curvature ^19, 23^, we repeated the mapping for four of these proteins in *h-* cells. Comparing the distance of each protein to their Myo52 reference showed nearly indistinguishable trajectories of Prm1, Fus1 and the exocyst subunit Exo84 in *h+* and *h-* cells (Figure S2A). The Myo51 trace was more distal (cell internal) in the *h-* cell but followed the same trend as in the *h+* cell. Therefore, the overall organization of the fusion focus is the same in both cell types. We thus extended the details of the map in *h+* cells only.

The distal localization of Myo51 was mirrored by that of tropomyosin Cdc8, which decorates linear actin filaments (Figure 2B) ^40^. The distal position of the bulk of Myo51-decorated actin filaments was confirmed using Airyscan super-resolution microscopy (Figure 2B, right). Thus, actin filaments initially extend a few hundred nanometers into the cytosol and then become shorter or more cross-linked as fusion proceeds. Compaction of the fusion focus also occurs in the lateral dimension along the cell-cell contact area (perpendicular to the fusion axis) ^19^. We had shown that Myo51 and Cdc8 contribute to this compaction ^28^, likely through the action of Myo51 sliding Cdc8-decorated actin filaments relative to each other ^27^. We confirmed this function by measuring the lateral distribution of Fus1 along the cell-cell contact area (perpendicular to the fusion axis), which was indeed wider in *myo51Δ* than WT cells (see below Figure 6A-B).

In the zone proximal to the partner cell, we located the transmembrane proteins Dni1 and Dni2, involved in membrane merging, in addition to Scd2 and Prm1 (Figure 2C) ^33, 34^. Although these four proteins have functions at the plasma membrane, their position did not overlap, with Scd2 exhibiting the trace most distant from Myo52. We reason that a pool of the transmembrane proteins (Prm1, Dni1 and Dni2) is also present on the secretory vesicles that deliver these proteins to the plasma membrane. As the cluster of secretory vesicles is at a distance from the plasma membrane (see below), the fluorescence centroid position of transmembrane proteins would be displaced internally in a manner proportional to the proportion of proteins associated with the vesicles. By contrast, Scd2, binds the plasma membrane peripherally by association with active Cdc42 GTPase ^41^. We attempted to localize the plasma membrane using the phosphatidylserine sensor probe LactC2 (Figure S2C-D). However, its trace was also internal, though parallel, to that of Scd2, likely for the same reason that LactC2 also binds phosphatidylserine on secretory vesicles ^42, 43^. Therefore, we used Scd2 as proxy for the position of the plasma membrane.

The medial zone between transmembrane proteins and actin filaments provided the most unexpected observations. The median position of the formin Fus1, which was initially slightly distal to that of Myo52, colocalized perfectly with Myo52 in the last 30 min before fusion (Figure 2B). By contrast, exocyst components Exo70, Exo84 and Sec6, as well as the secretory vesicle-associated Rab GTPases Ypt2 (homologous to Rab8/Sec4 in *S. cerevisiae*) and Ypt3 (homologous to Rab11) ^44, 45^, all initially localized close to Myo52. The overlap in position is particularly clear when we examine traces at even earlier times than 60 min before fusion (Figure S2B). However, they then appeared to move distally away from Myo52 up to fusion time (Figure 2D). Exocyst and Rab GTPase traces did not perfectly overlap. Notably, Sec6 was more membrane-proximal, possibly because a pool of this protein at the plasma membrane induces displacement of the centroid. However, we conclude from their parallel traces that these describe the behavior of secretory vesicles. Thus, in the hour that precedes cell fusion, as the focus compacts, Myo52 detaches from the bulk of secretory vesicles and colocalizes with Fus1.

Because we used Myo52 as reference, its position appears static while other proteins move around it. Interpretation would be easier with the plasma membrane position as static reference. For easier visualization of the evolution of the focus over time, we used an idealized (smoothened) Scd2 trace as plasma membrane position, shifted the y-axis origin to this new reference point, and computationally transposed all other trajectories, yielding the map shown in Figure 2E. This visualization clearly illustrates our main findings: 1) the ensemble of proteins at the focus is progressively more compact and membrane-proximal, with all components moving towards the plasma membrane over time, with the notable exception of secretory vesicles, 2) the pool of secretory vesicles occupies a position at constant ∼ 100 nm distance from the plasma membrane, 3) the myosin V Myo52 detaches from vesicles and colocalizes with the formin Fus1 at an intermediate position.

### Two distinct functions for the myosin V Myo52

The observation that Myo52 and secretory vesicles occupy progressively distinct localizations in the fusion focus raises questions about the role of Myo52. A large body of previous work, mainly on Myo52’s orthologue Myo2 in *S. cerevisiae* during polarized cell growth, has shown that the motor associates with secretory vesicles from their budding at the trans-Golgi network all the way to their arrival at the plasma membrane, from where the myosin detaches only 2 to 3 seconds before vesicle fusion ^2, 12, 13^. Our observation that the bulk of Myo52 leaves vesicles in the fusion focus suggests otherwise during the process of cell-cell fusion.

To gain further insight, we analyzed the evolution of the local fluorescence intensity of the proteins described above, which is directly proportional to the number of molecules in the focus. We observed two main types of recruitment dynamics. The signal intensity of most proteins increased, peaked at fusion time and sharply dropped post-fusion. This increase in fluorescence as the focus becomes more compact indicates it also becomes denser. This was the case for Myo52 (as in previous manual measurements; ^19^), actin regulators (Fus1, Cdc8, Myo51), transmembrane proteins (Prm1, Dni1, Dni2) and polarity factors (Scd2), although the slope of their accumulation varied (Figure S3A). By contrast, secretory vesicle markers (Exo70, Exo84, Sec6, Ypt2, Ypt3) showed more moderate increase and reached a plateau about 40 min pre-fusion before decreasing 10-20 min before fusion (Figure S3B). A comparison of a subset of these factors in *h+* and *h-* cells showed that fluorescence intensity dynamics were similar when measured in either cell type, but recruitment seemed to be slightly delayed in *h-* cells (Figure S3C). A direct comparison of the recruitment dynamics of Myo52, Fus1 and secretory vesicles, extended to 100 min before fusion, shows parallel accumulation of vesicles and the myosin up to 40 min before fusion, when vesicular markers reach a plateau (Figure 3A). Myo52 then continues to accumulate, coincident with strong accumulation of Fus1. These observations led us to hypothesize that Myo52 plays two distinct consecutive roles: it first transports secretory vesicles; it then plays a second role in organizing the focus alongside Fus1. We set out to test this hypothesis.

**Figure 3:**
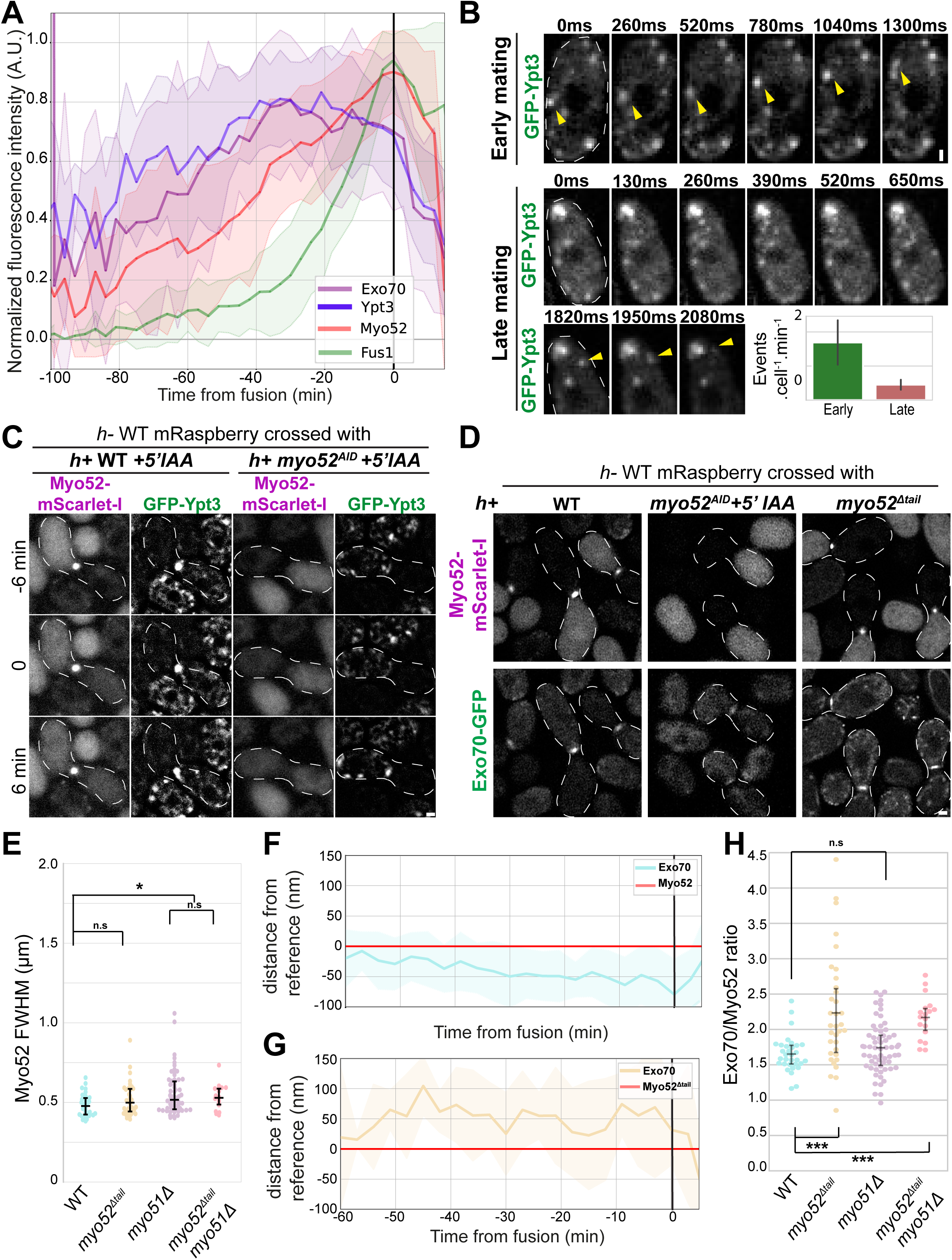
Myo52 concentrates secretory vesicles on the fusion focus. **A.** Fluorescence intensity of Myo52, Exo70, Ypt3 and Fus1 at the fusion focus during the fusion process. **B.** Spinning disk timelapse images of Ypt3 linear movements during early mating and cell-cell fusion (late mating). Yellow arrowheads track individual vesicles. The graph shows the average number of linear movements observed per cell per minute over at least 4 consecutive time points. **C.** Deltavision timelapse images of GFP-Ypt3 and Myo52-mScarlet-I in WT or Myo52^AID^-mScarlet-I in *myo52^AID^ h+* cells mated with *h-* WT cells expressing mRaspberry before and after cell fusion. t0 corresponds to entry of the mRaspberry signal in the *h+* cell. The cells were treated with 100 nM 5-adamantyl-IAA (5’IAA) during starvation for 5h. Note the complete depletion of Myo52^AID^-mScarlet-I. **D.** Airyscan images of Myo52-mScarlet-I and Exo70-GFP in indicated pre-fusion *h+* strains mated with *h-* WT cells expressing mRaspberry and Myo52-mTagBFP2 (mTagBFP2 signal not shown). The mutants Myo52 alleles were also tagged with mScarlet-I. **E.** FWHM of Myo52-mScarlet-I fluorescence profiles perpendicular to fusion axis in cells as in (D) and indicated *myo51Δ* cells at the last time point before fusion or the last time point when the WT cell Myo52-mTagBFP2 signal was round. **F-G.** Median directed distances of Exo70 from Myo52 (F) and Myo52^Δtail^ (G). **H.** Ratio between Exo70 FWHM and Myo52 FWHM from profiles as in (E) measured in the same cell. On all graphs, shaded zones show standard deviation, black line shows fusion time. Scale bars are 1 µm.

### Myo52 transports secretory vesicles in preparation for fusion

We first examined whether Myo52 indeed transports secretory vesicles. During mitotic growth, we readily observed long-distance linear movements of coincident Myo52-mScarlet-I and GFP-Ypt3 signals, in agreement with previous observations (Figure S4A). Long-range movements (defined as spanning at least 4 consecutives timepoints) were observed on average 7 times per cell per min (Figure S4B). During sexual differentiation, Myo52-mScarlet-I and GFP-Ypt3 signals were weaker and photobleached more rapidly, preventing fast two-color imaging. We thus acquired single-channel time lapses, to assess and quantify long-range movements of GFP-Ypt3. We observed long-range movements during the exploratory phase of mating, when unstable polarity patches probe cells’ environment to find a partner ^39, 46^, at a frequency of around 2 events per cell per min (Figure 3B, Movie S1). In cells with a stable fusion focus, identified by focal accumulation of Ypt3, time lapse imaging gave a sense of directed vesicle movement towards the focus (Movie S2), but at close range and rarely over > 3 time points (Figure 3B). Thus, secretory vesicles are transported vectorially, though this activity may be reduced during fusion.

To test more directly whether Myo52-based transport is required to bring vesicles to the fusion focus, we used an auxin-inducible degron allele of *myo52* (*myo52^AID^*), which allowed to decouple a role during cell fusion from prior cell polarization defects. Upon depletion of Myo52, Ypt3 did not accumulate at the fusion focus (Figure 3C). A thin, wide localization at the fusion site was occasionally observed (Figure 3C, Movies S3). Upon cell fusion with a WT partner, Ypt3 was immediately observed at the fusion site, showing that secretory vesicles in the mutant cells can be captured by the WT Myo52 (Figure 3C, Movie S3). Thus, Myo52 is required to concentrate secretory vesicles on the focus, as previously observed for their glucanase cargoes ^19^.

We noted that, in contrast to Ypt3, Exo70 still localized at the fusion site, albeit weakly, in *myo52^AID^* cells (Figure 3D). This is reminiscent of our observations during mitotic growth where the exocyst complex normally associates with secretory vesicles ^11^, but in absence of actin-based transport still localizes to cell poles, where it is able to capture vesicles for polarized growth ^47^. To probe this point further, we used a mutant allele of Myo52 lacking its C-terminal cargo-binding domain, which mediates association with secretory vesicles ^7^. This truncated Myo52^Δtail^ protein remains competent for actin-based movements, as it localizes to the poles of vegetative cells ^18, 48^ (Figure S4C) and accumulates at the fusion focus in mating cells, where its lateral distribution (described below as signal width) is not significantly different from that of full-length Myo52 (Figure 3D-E). Thus, the organization of the actin fusion focus is not grossly altered in *myo52^Δtail^* mutant cells, even though these mutants fail to concentrate cell wall digestive enzymes and are largely fusion deficient ^19^. In *myo52^Δtail^* cells, like in *myo52^AID^*, the exocyst subunit Exo70-GFP remained localized to cell poles during mitotic growth (Figure S4C) and localized to the fusion site in mating cells (Figure 3D). However, both the position of its centroid and the width of its distribution were distinct from that in WT cells. In WT cells, Exo70 signal was detected at a cell-internal position relative to Myo52, as described above (Figure 3F). By contrast, in *myo52^Δtail^* cells, Exo70 signal was closer to the plasma membrane than the myosin (Figure 3G). The width of the Exo70 signal, which is ∼ 1.6-fold larger than that of Myo52 in WT cells – reflecting the larger area occupied by the vesicle pool – was even wider in *myo52^Δtail^* (Figure 3H). We note that deletion of the actin crosslinking myosin V Myo51 had a small effect on Myo52 distribution (Figure 3E), as reported ^28^, but did not affect the relative distribution of Exo70 and Myo52 (Figure 3H). The membrane-proximal and wider localization of Exo70 in *myo52^Δtail^* leads us to conclude that, whereas the exocyst primarily decorates secretory vesicles on the focus in WT cells, it localizes to the plasma membrane in absence of functional Myo52.

In summary, binding of the myosin V Myo52 to secretory vesicles promotes their early transport to the fusion site and their concentration on the fusion focus. In absence of Myo52, vesicles are likely captured at the plasma membrane by the exocyst complex.

### Myo52 colocalizes and interacts with the formin Fus1

We examined the relationship between Myo52 and Fus1. In addition to their colocalization along the fusion axis (Figure 2B), Fus1 and Myo52 exhibited almost identical signal widths, with Myo52 slightly more compact than Fus1 (Figure 4A-C). In *fus1Δ* cells, Myo52 did not form a compact dot and instead decorated a wide region at the fusion site (Figure 4A) ^19^. Similarly, in cells in which Fus1 is present but fails to condense due to deletion of its intrinsically disordered domain (*fus1^ΔIDR^*) ^26^, Fus1 and Myo52 exhibited similarly wider distributions (Figure 4B-C). Thus, Myo52 distribution follows that of Fus1, or the actin filaments it assembles.

**Figure 4:**
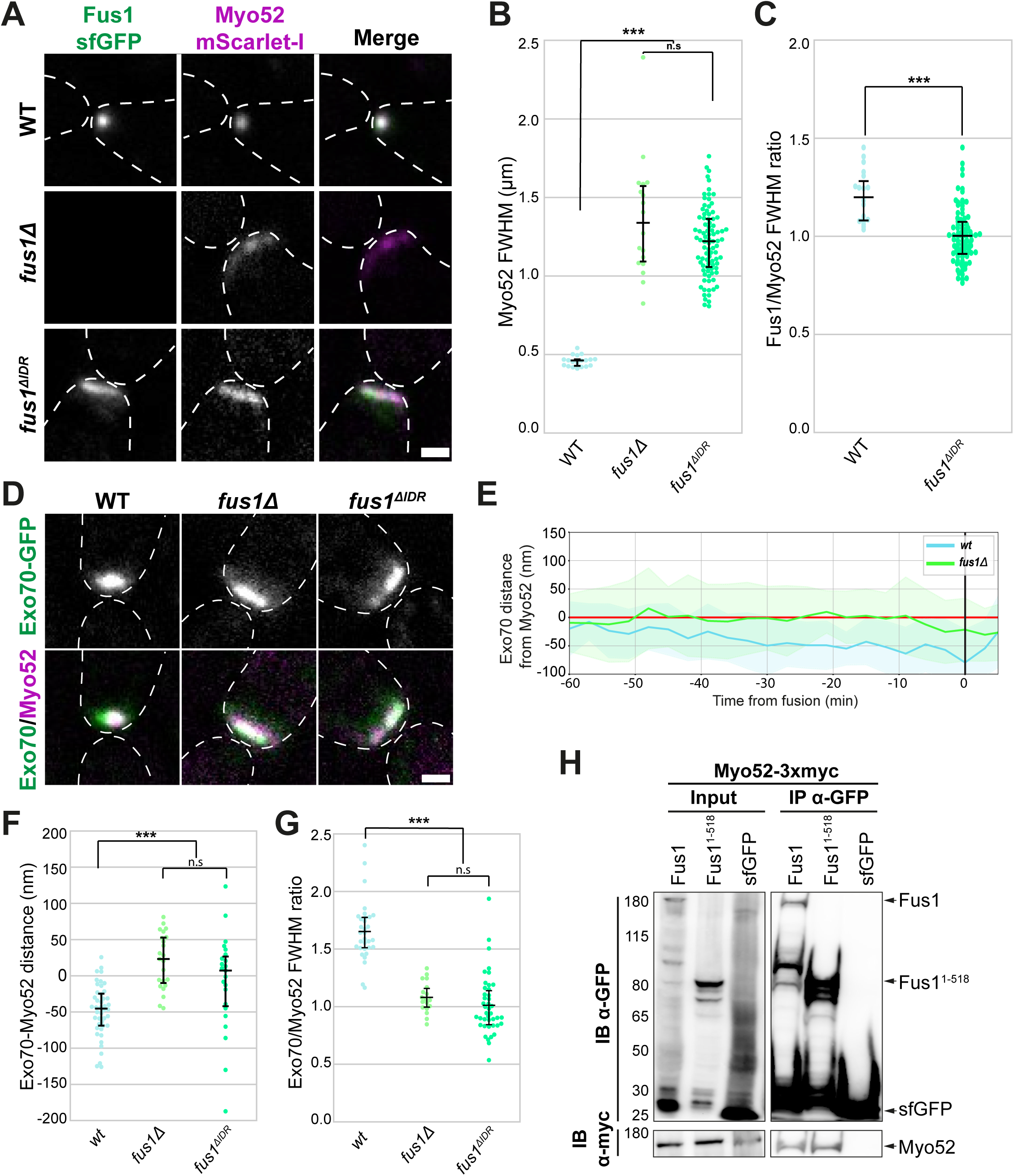
Myo52 interacts with Fus1. **A.** Images of Fus1-sfGFP and Myo52-mScarlet-I in WT (top), *fus1Δ* (middle) and *fus1^ΔIDR^ h+* cells mated with WT *h-* cells (Myo52-mTagBFP2 signal not shown). **B.** FWHM of Myo52-mScarlet-I fluorescence profiles perpendicular to fusion axis in cells as in (A) at the last time point before fusion or the last time point when the WT cell Myo52-mTagBFP2 signal was round. **C.** Ratio between Myo52 FWHM and Fus1 FWHM from profiles as in (B) measured in the same cell. **D.** GFP fluorescence and merge images of Exo70-GFP and Myo52-mScarlet-I in WT, *fus1Δ* and *fus1^ΔIDR^ h+* cells. **E.** Median directed distances of Exo70 from Myo52 in WT and *fus1Δ h+* cells over the fusion process. **F.** Median directed distances of Exo70 from Myo52 in WT, *fus1Δ* and *fus1^ΔIDR^ h+* cells in unsynchronized cells. **G.** Ratio between Exo70 FWHM and Myo52 FWHM from profiles as in (D) measured in the same cell. **H.** Co-immunoprecipitation of Myo52-3xmyc with Fus1-sfGFP and Fus1^1–518^–sfGFP expressed in vegetative cells under *pnmt1* promoter. sfGFP serves as negative control. Scale bars are 1 µm.

Interestingly, in *fus1Δ* and *fus1^ΔIDR^*, the position of secretory vesicles, marked by Exo70, along the fusion axis coincided with that of Myo52 during the entire fusion process (Figure 4D-F) and their distribution was as wide as that of Myo52 (Figure 4G). We note that in these mutants, we could not use automated centroid tracking due to the wide distribution of the Myo52 reference and instead defined the fusion axis manually, which we verified yields similar results (Figure S1C, method 3). Because very few *fus1^ΔIDR^* mutant cells successfully fuse, even with a WT partner, we could not obtain measurements over the whole fusion time course in this mutant. We instead measured the average distance between Exo70 and Myo52 at random times before cell fusion, which gave a ∼ 50 nm distance in WT, but ∼ 0 nm in both *fus1^Δ^*^IDR^ and fus1*Δ* (Figure 4F). We conclude that the distal position of vesicles in WT cells is a steric consequence of the architecture of the fusion focus, which restricts space at the center of the actin aster and promotes separation of Myo52 from the vesicles.

These observations led us to test whether Myo52 physically associates with Fus1. To this aim, we conducted co-immunoprecipitation experiments from mitotic cells in which tagged Fus1 and Myo52 were co-expressed (Figure 4H). Myo52 co-immunoprecipitated with full-length Fus1. Importantly, it also co-immunoprecipitated with Fus1 N-terminus (aa 1-518), which lacks the IDR and the FH1-FH2 actin assembly domains, indicating that the interaction is not mediated by Myo52 being caught on Fus1-nucleated actin filaments. Recent work in *S. cerevisiae* showed an interaction between the Myo52 homologue and Pea2, a component of the polarisome, a multicomponent complex that assembles around the protein Spa2 and regulates the formin Bni1 function ^49^. However, Pea2 is not conserved outside close relatives of *S. cerevisiae* ^50^. We also found that Spa2 does not concentrate at the fusion focus in *S. pombe* but decorates a wide area at the plasma membrane (Figure S5). Thus, the polarisome is unlikely to serve as intermediate between Fus1 and Myo52. We conclude that Myo52 directly or indirectly binds the formin Fus1, independently of actin assembly.

### Myo52 transports Fus1

The colocalization and interaction of Fus1 with Myo52 suggest that Fus1 may be a Myo52 cargo. In previous work, we found that Fus1 IDR can be functionally fully replaced by the low-complexity region of Fused-in-Sarcoma (FUS^LC^) ^26^. In serendipitous observations of chimeric Fus1^ΔIDR^-FUS^LC^, using a FUS^LC^ variant with 27 additional arginine residues (FUS^LC-27R^) that forms solid aggregates ^51^, we unexpectedly observed expression and formation of small Fus1^ΔIDR^-FUS^LC-27R^ foci in mitotically growing cells. While the underlying cause of the mitotic expression/stabilization of this chimeric protein is unknown, by clustering Fus1 outside its physiological context, this variant offered an opportunity to investigate its possible transport by Myo52.

Fus1^ΔIDR^-FUS^LC-27R^ foci showed extensive colocalization with Myo52 and both proteins exhibited co-linear long-range movements (Figure 5A, movies S4-S5). We quantified the number of long-range movements, defined as continuous movements crossing at least one quarter of the cell without a pause, in co-labelled cells imaged at 600 ms interval over a minute. Treatment with Latrunculin A, which leads to F-actin depolymerization and Myo52 dispersion, blocked all such movements (Figure 5B), indicating that they are actin-dependent.

**Figure 5:**
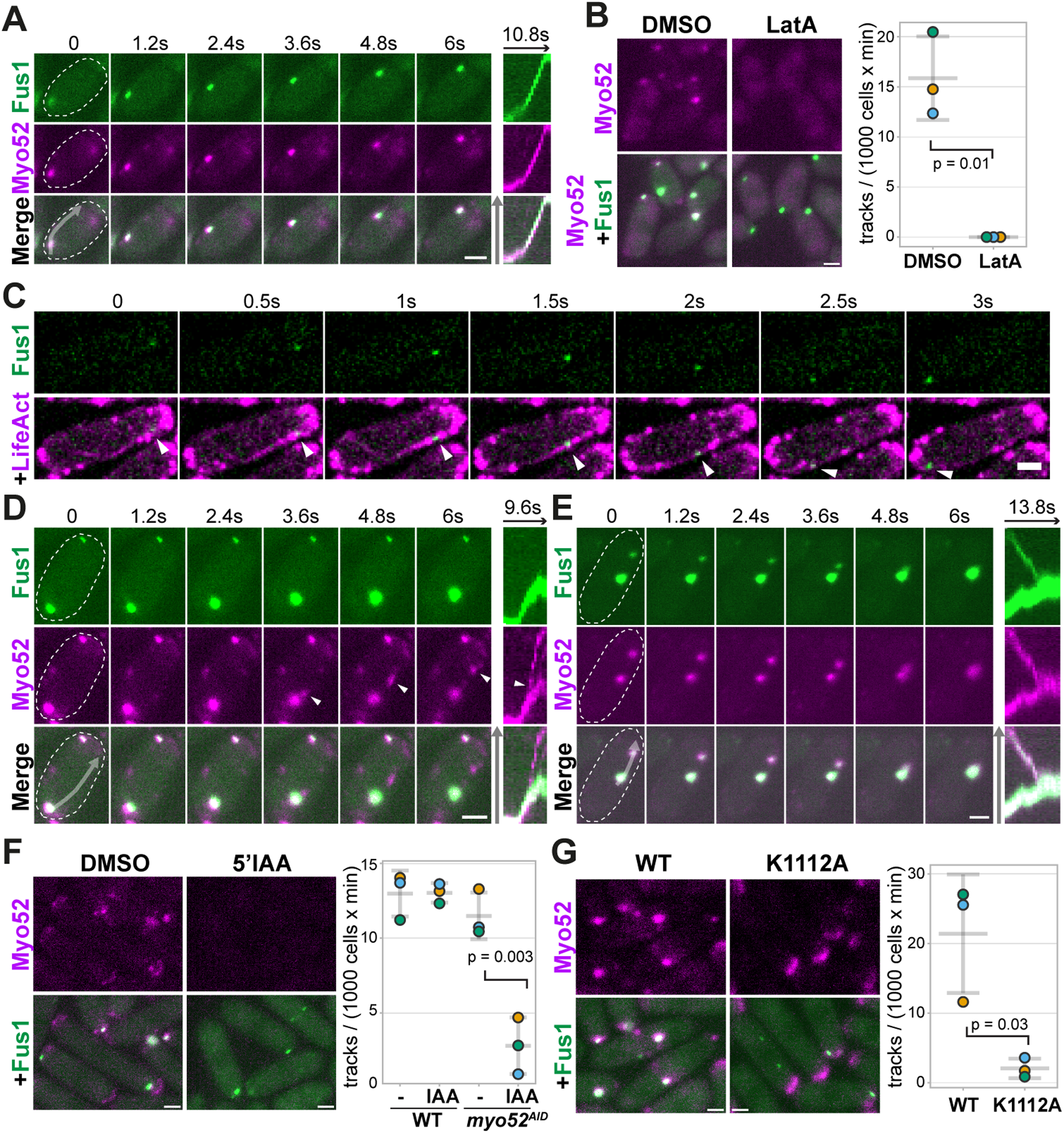
Myo52 transports Fus1. **A.** Timelapse images of Fus1^ΔIDR^-FUS^LC-27R^-sfGFP and Myo52-mScarlet-I in vegetative cells. The kymograph on the right was constructed along a segmented line along the trajectory of the coincident Fus1-Myo52 signal (grey arrow). **B.** Cells expressing Fus1^ΔIDR^-FUS^LC-27R^-sfGFP and Myo52-mScarlet-I were mounted on 250μl MSL-N agar imaging pad with 1μl of 20mM LatA (final concentration 80μM) or 1μl of DMSO solvent. The graph indicates the frequency of long-range tracks observed in 3 independent experiments. N > 1000 cells for each experiment. **C.** Spinning disk timelapse images of Fus1^ΔIDR^-FUS^LC-27R^-sfGFP and LifeAct-mCherry in vegetative cells. **D-E.** Timelapse images of Fus1^ΔIDR^-FUS^LC-27R^-sfGFP and Myo52-mScarlet-I in vegetative cells showing continued movement of Myo52 without Fus1 (D; arrowheads) and convergent movements (E). The kymographs on the right were constructed along a segmented line along the trajectory of the coincident Fus1-Myo52 signal (grey arrows). **F.** *myo52^AID^* cells expressing Fus1^ΔIDR^-FUS^LC-27R^-sfGFP and Myo52-mScarlet-I treated or not with 100 nM 5’IAA. The graph indicates the frequency of long-range tracks observed in 3 independent experiments in these cells and WT (*myo52+*) controls. N > 1000 cells for each experiment. **G.** Fus1^ΔIDR^-FUS^LC-27R^-sfGFP with WT (left) or K1112A mutant FH2 domain (right) and Myo52-mScarlet-I in vegetative cells. The graph indicates the frequency of long-range tracks observed in 3 independent experiments. N > 1000 cells for each experiment. Scale bars are 2 µm.

**Figure 6:**
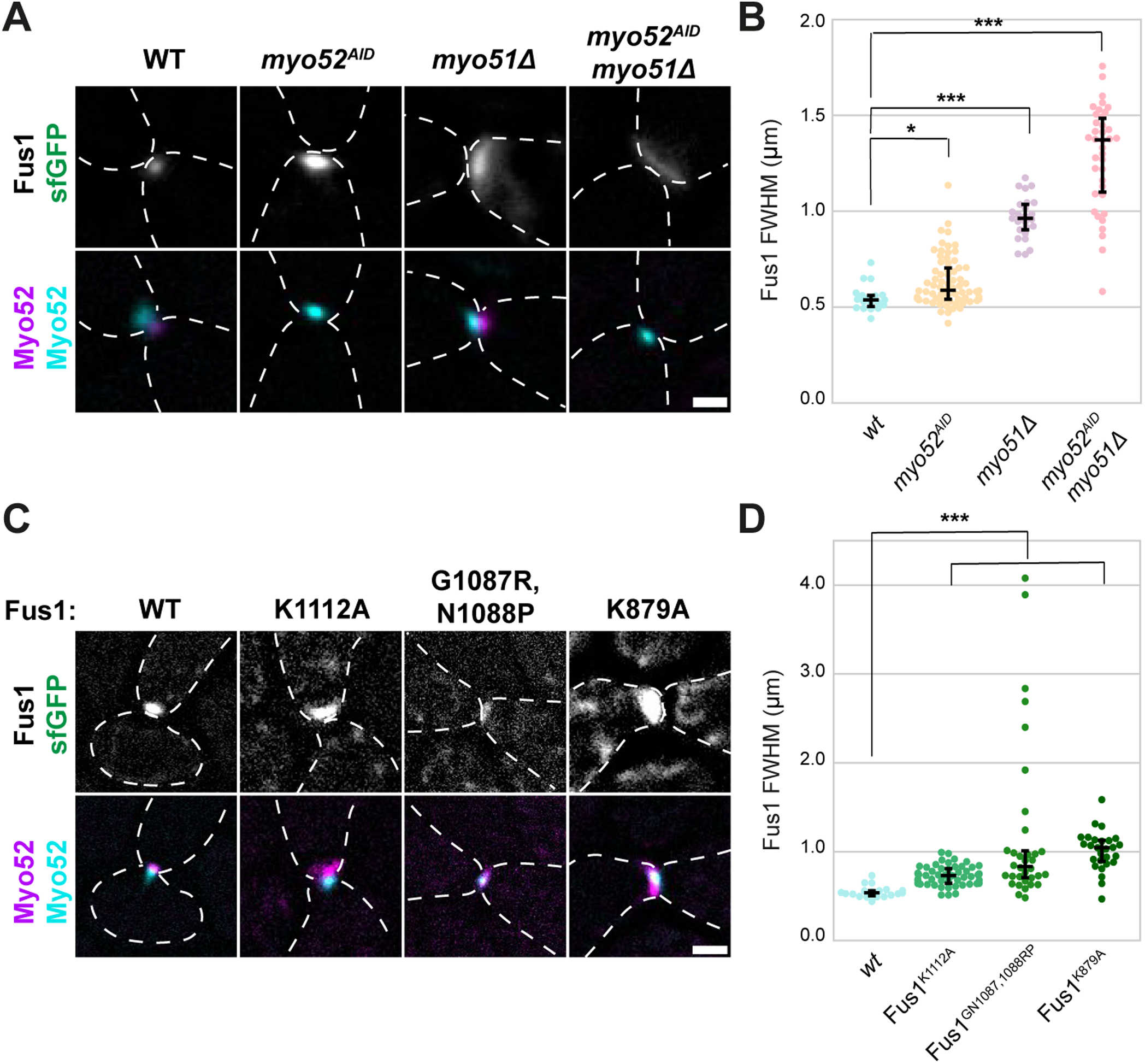
Myo52 and Fus1 activity are required for actin focus compaction. **A.** Fus1-sfGFP (top) in WT, *myo52^AID^*, *myo51Δ* and double mutant *h+* cells. The *myo52^AID^* mutant was treated with 100nM 5-adamantyl-IAA (5’IAA) for 5h. The bottom row shows the Myo52-mScarlet-I signal in the same cell and the Myo52-mTagBFP2 signal in the partner WT cell. **B.** FWHM of Fus1-sfGFP fluorescence profiles perpendicular to fusion axis in cells as in (A) at the last time point before fusion or the last time point when the Myo52-mTagBFP2 signal was round. **C.** Fus1-sfGFP in *h+* cells expressing Fus1 that is either WT or carrying the indicated point mutations in its FH2 domain. The bottom row shows the Myo52-mScarlet-I signal in the same cell and the Myo52-mTagBFP2 signal in the partner WT cell. **D.** FWHM of Fus1-sfGFP fluorescence profiles perpendicular to fusion axis in cells as in (C) at the last time point before fusion or the last time point when the Myo52-mTagBFP2 signal was round. Scale bars are 1 µm.

Several observations suggested that the Fus1 foci move along pre-existing actin cables nucleated by either other Fus1, or the endogenous formins For3 or Cdc12 expressed during mitotic growth. First, co-imaging of Fus1^ΔIDR^-FUS^LC-27R^ and actin cables labelled with LifeAct-mCherry showed foci moving along actin cables (Figure 5C, Movie S6). Second, we observed instances of divergent Fus1^ΔIDR^-FUS^LC-27R^ and Myo52 movements where the two proteins were initially coincident, but the formin movement stopped and a Myo52 dot appeared to continue along the same initial trajectory (Figure 5D, Movies S7-S8). This indicates the existence of a continuous underlying track spanning ahead of the Fus1 focus. Third, we observed instances of convergent Fus1^ΔIDR^-FUS^LC-27R^ and Myo52 movements, leading to the apparent fusion of two Fus1^ΔIDR^-FUS^LC-27R^ foci, suggesting that actin-based movements of Fus1 contribute to its clustering (Figure 5E, Movie S9-S10).

To directly test whether Myo52 powers Fus1^ΔIDR^-FUS^LC-27R^ long-range movements, we used the *myo52^AID^* mutant to deplete Myo52 from the cell. Quantification of the frequency of long-range movements in repeated experiments showed a strong reduction after Myo52 depletion (Figure 5F), indicating that Myo52 is required for many of the observed movements. The remaining long-range movements may be due to Fus1-dependent actin polymerization powering its own movements, as has been observed for some formins in other organisms ^52^. We aimed to test for this possibility by introducing the K1112A mutation in the FH2 domain of Fus1^ΔIDR^-FUS^LC-27R^ previously shown to abolish actin assembly in vitro and reduce function in vivo ^30, 53^. This mutant showed very few long-range movements (Figure 5G). However, Myo52 also failed to localize on the mutant Fus1 foci, which indicates that the enrichment of Myo52 on Fus1 foci requires Fus1 to assemble actin filaments. This may help increase Myo52 concentration in the vicinity of Fus1, favoring their encounter. Thus, even though Myo52 can bind Fus1 independently of actin assembly, as shown by the association of Fus1 N-terminus with Myo52 (Figure 4H), efficient capture of Fus1 by Myo52 may be promoted by Fus1 activity. Therefore, the K1112A mutant may not clearly distinguish Fus1-powered from Myo52-powered movements.

Together, the association of Fus1 with Myo52, their overlapping localizations and the Myo52-dependent long-range movement of Fus1 demonstrate that the formin is a cargo of the myosin V.

### Myo52 and Fus1 activity focalize the fusion focus

The identification of Fus1 as a Myo52 cargo suggest a focusing mechanism by which Myo52 transports Fus1 along filaments nucleated by other Fus1 dimers, leading to focusing of Fus1 and actin filament barbed ends at the fusion focus. This model predicts that both Myo52-based transport and Fus1 actin assembly activity contribute to setting the dimensions of the fusion focus.

To test the role of Myo52 in modulating the localization of Fus1 at the fusion site, we used the *myo52^AID^* mutant and measured the width of the Fus1 signal in cells paired with a WT partner, in which we could assess Myo52 focus formation, to ensure that any phenotype was not amplified by problems in cell-cell signaling ^24^. Upon Myo52^AID^ depletion, Fus1 occupied a wider region than in WT cells (Figure 6A-B), in agreement with previous measurement of Fus1 in *myo52Δ* cells (in which both partner cells were mutant) ^19^. Myo52 depletion also led to a wider Fus1 distribution in a *myo51Δ* mutant background, indicating both myosins contribute to focus condensation through distinct pathways. We note that depletion of either *myo52* or *myo51* did not alter the wide distribution of Fus1^ΔIDR^ (Figure S6A-B).

To test on which filaments Myo52 transports Fus1, we measured Fus1 focus dimensions in *for3Δ* and in *fus1* mutants with reduced actin assembly activity. In *for3Δ*, cell fusion was delayed, but Fus1 formed a focus of normal dimensions (Figure S6C-D), indicating that For3-nucleated actin cables are not essential for formation of the fusion focus. By contrast, three distinct point mutants in Fus1 FH2 domain, previously shown to block actin assembly in vitro and to exhibit reduced functionality in vivo ^30, 53^, led to a wider distribution of Fus1 at the fusion site (Figure 6C-D). We conclude that concentration of the formin Fus1 to form the fusion focus requires its transport by Myo52 along the filaments it assembles.

## Discussion

In this work, we present a high-resolution spatiotemporal map of the actin fusion focus in the fission yeast *Schizosaccharomyces pombe*. Key features of the fusion focus are the membrane-proximal position of the formin Fus1 condensate, from where tropomyosin-decorated actin filaments extend, and a more distal pool of secretory vesicles. The map reveals that the fusion focus compacts over time, with most components accumulating and becoming more plasma membrane-proximal until fusion pore opening. Secretory vesicles are an exception to this rule: they reach a plateau ∼ 40 min before fusion and stay at constant ∼ 100 nm distance from the plasma membrane throughout the process. We demonstrate that the myosin V Myo52, which initially transports vesicles to the focus, dissociates from the vesicle pool and joins the formin Fus1 (Figure 7). By binding and transporting the formin along the filaments it nucleates, it underlies positive feedback that ensures convergence of the formin and associated actin filaments in a tight focus.

**Figure 7:**
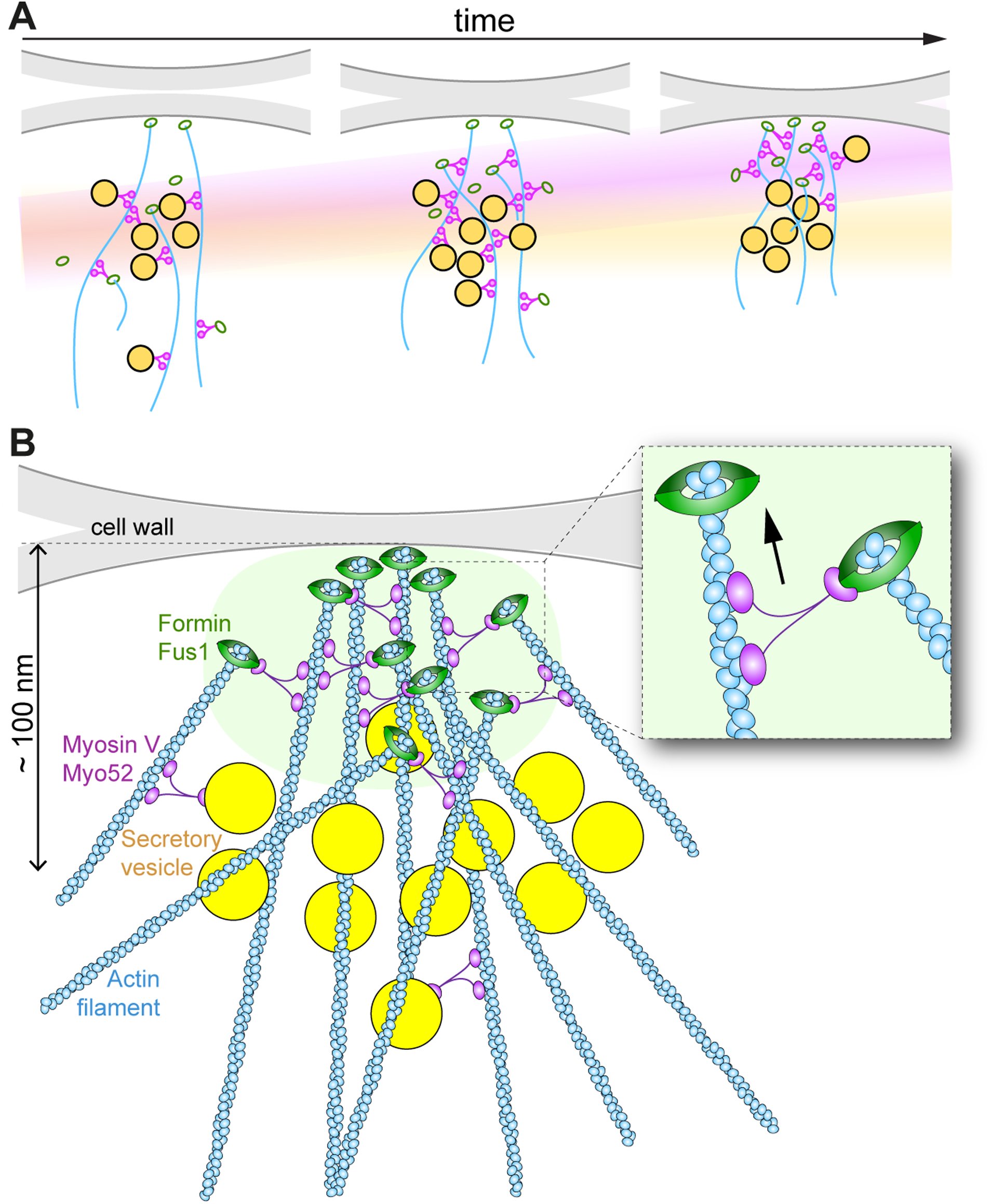
Schematic model of the actin fusion focus. **A.** Schematics of the fusion focus over time, showing Myo52 transporting both secretory vesicles and Fus1. The transport of Fus1 along filaments assembled by other Fus1 dimers promotes the convergence of Fus1 and actin barbed ends together with Myo52, whose centroid is moving closer to the plasma membrane over time (magenta haze). Myo52 also transports secretory vesicles, but their large size prevents their transport to the center of the aster, leading to a reserve pool at constant distance from the plasma membrane (yellow haze). **B.** Focusing of Fus1 and the actin aster requires Fus1 biomolecular condensate formation (green shaded area), transport by Myo52 (inset) and actin crosslinking activities (not shown). Vesicles are ∼ 60 nm in diameter and drawn at size smaller than reality.

### Fluorescence centroid mapping

The fluorescence centroid tracking approach we have developed provides high precision – measured at ∼ 8 nm from doubly-tagged proteins – while allowing live acquisition with a simple epifluorescence microscope. At the start of the work, we assumed that proteins’ relative spatial position and order of events would be reproducible from cell to cell, an assumption validated by the finding that distinct proteins exhibit distinct profiles. The map derives precision from this stereotyped behaviour and the median position of proteins can be viewed as an idealized fusion focus. In principle, the method can be used with appropriate spatial and temporal reference points and standard equipment to map any event occurring in a stereotypical manner. For easy re-use and adaptation, all codes are available from Github.

While the median position is extremely precise, it is important to understand that there may be substantial variations in the position of individual proteins at any given time. For a given protein, if all individual molecules behave the same, the centroid position will accurately reflect its average localization. If the vast majority behave in one way and a few have distinct localization, the centroid position will likely not be altered by the few and the map will reflect the localization of the majority. For instance, even if the Myo52 profile colocalizes perfectly with that of Fus1, it does not exclude that a few molecules of Myo52 transport a few vesicles from the cytosol into the focus or towards the plasma membrane. Finally, if a protein forms two pools located at a distance lower than optical resolution, then the median position will represent a weighted average of these two positions. This is likely the case for transmembrane proteins, which must localize to the plasma membrane, as this is their site of action, but are transported there by secretory vesicles.

### Organization of vesicles and the exocyst in the fusion focus

Our map demonstrates that secretory vesicles, decorated by the Rab GTPases Ypt3 (Rab11 homolog) and Ypt2 (Sec4/Rab8 homolog), both marking secretory vesicles in a wide range of eukaryotes, accumulate at the fusion site early in the fusion process, where they remain at constant ∼100 nm average distance from the plasma membrane. These findings are consistent with ultrastructural analysis, which showed constant density of 60 nm diameter secretory vesicles throughout the process, except at very early stages ^23^. The vesicles are also decorated by the exocyst complex, whose profiles have similar slopes and are nearly overlapping with those of Rab GTPases, with the possible exception of Sec6 (see below). While initially coincident with the myosin V Myo52, in the 30-40 min before fusion, vesicles are no longer bound by Myo52. We considered the possibility that the displacement between Myo52 and Rab GTPases traces may arise from localization of Myo52 to the front of the vesicles relative to a more uniform peri-vesicular Rab GTPase distribution. However, this would result in a smaller displacement equivalent to the vesicle radius (30 nm, whereas we observed ∼ 60 nm). Thus, the pool of vesicles is ready for secretion, but not yet tethered at the plasma membrane. The distal location of this vesicle pool, relative to Fus1 formin, is likely a consequence of the steric barrier imposed by the convergence of actin filament barbed ends in the fusion focus.

It is interesting to compare the organization of the exocyst complex in the focus to that in other species. The hetero-octameric exocyst complex serves to tether secretory vesicles at the plasma membrane through redundant interactions of the subunits Exo70 and Sec3 with phosphoinositides and small Rho-family GTPases. Early work in *S. cerevisiae* proposed that these two subunits act as landmarks at the plasma membrane ^54, 55^. However, subsequent studies showed that localization of Sec3 like other components requires actin cables and myosin V-mediated transport ^56^, failed to observe subcomplexes ^57, 58^, and visualized all eight subunits on travelling vesicles ^2^. In *S. pombe*, exocyst subunits, including Sec3 and Exo70, are present on secretory vesicles during mitotic growth ^11^ and, as we show here, during cell-cell fusion. We note however that subcomplexes have been observed in mammalian cells ^59^, and that in *Neurospora crassa* hyphae, different exocyst subunits decorate the vesicle-rich Spitzenkörper (Exo70/Exo84) and the plasma membrane (other subunits including Sec6) ^60^. Our observation that the Sec6 profile is slightly displaced towards the plasma membrane suggests that distinct subcomplexes may also exist in *S. pombe* and that, like in *Neurospora*, a pool of Sec6 may decorate the plasma membrane, perhaps through its known interaction with the t-SNARE ^61^, offsetting the median fluorescence position.

Contrary to the situation in budding yeast, in *S. pombe*, the localization of the exocyst to the poles of mitotic cells does not require actin-based transport ^47, 62^. Similarly, the exocyst can localize to the fusion site without actin-based transport, but we now show that its precise position is distinct: in *myo52^Δtail^*, Exo70, which localizes distally with the vesicle pool in WT cells, now accumulates proximally and over a wider zone at the plasma membrane. Because polarized growth is not abrogated in *myo52* mutant cells and we observe a thin, wide Ypt3 signal at the fusion site, the exocyst at the plasma membrane is likely able to capture secretory vesicles, which however do not transit through the fusion focus. Our data demonstrate that Myo52 plays a key role in recruiting secretory vesicles to the fusion focus, ensuring that they do not reach the plasma membrane at random location. It is tempting to speculate that Myo52 also promotes exocyst complex recruitment on the vesicle, for instance through the described conserved interface on Sec15 ^6^ to ensure vesicle funneling through the actin fusion focus.

### Compaction of the fusion focus

One major finding from our precise map is the movement of Myo52 away from the vesicle pool and its colocalization with the Fus1 formin. The proximity of the two proteins, their physical association and the observation that Fus1 clusters are moved along actin cables by the myosin in mitotic cells establish a functional interaction between the two proteins. Because formation of a tight Fus1 focus requires both Myo52 and the actin assembly function of Fus1 but not For3-nucleated actin filaments, this supports the view that Myo52 transports Fus1 along actin filaments nucleated by other Fus1 dimers, leading to focusing of Fus1-decorated actin filament barbed ends (Figure 7).

Our previous work showed that Fus1 IDR is absolutely essential for focus formation and can be replaced by heterologous phase-separating protein domains, indicating that the fusion focus forms through biomolecular condensation ^26^. In this context, and describing the focus in biophysical terms, Myo52-dependent Fus1 transport promotes cohesive forces inside the condensate, increasing condensate surface tension to form a more spherical structure. This combines with actin cross-linking functions of the structural myosin V Myo51 ^28^ and likely of the Fus1 FH2 domain itself, which we did not re-examine here ^21^. In absence of these cohesive forces, the weak multivalent Fus1 self-interactions are insufficient, leading to a condensate that wets the plasma membrane and thus adopts an elongated shape.

The conjunction of self-organization principles from cytoskeletal polymer biology and biomolecular condensation makes the fusion focus a very interesting structure for future investigations and raises important questions. Notably, as vesicles remain largely outside of the Fus1 condensate and the bulk of Myo52 dissociates from them, one critical question is how they cross the Fus1 condensate to reach the plasma membrane. During budding in *S. cerevisiae*, the type V myosin remains vesicle-bound until vesicle tethering at the plasma membrane ^2^. While this is clearly not the case in the fusion focus, it is possible that Myo52 re-associates with individual vesicles as they travel the remaining distance to the plasma membrane. Another outstanding question is whether the Fus1-loaded Myo52 preferentially recognizes Fus1 rather than For3-nucleated filaments. This should be addressed in future work.

### Commonality and diversity in cytoskeletal aster formation

The mechanisms of actin fusion focus formation are highly reminiscent of those underlying the organization of spindle poles, where microtubule minus ends converge (reviewed in ^63^). Notably, formation of a microtubule aster from the centrosome relies on principles of biomolecular condensation ^64^ and on the transport of filament ends by motors. Spindle pole focusing is largely dependent on the dynein-dynactin-NuMA complex ^65–67^, where NuMA binds the MT minus ends and recruits the dynein-dynactin motor complex ^68^. In turn, the dynein-dynactin complex transports the bound microtubule end towards the minus end of other microtubules. Thus, forming a cytoskeletal aster, whether microtubule or actin-based, relies on common principles.

The fusion focus bears close similarity with the Spitzenkörper (‘tip body’), an accumulation of secretory vesicles proposed to act as supply center for fast hyphal growth in filamentous fungi ^69^. In *Aspergillus nidulans*, the sole formin SepA localizes as a tight dot at the hyphal tip, and, like Fus1, requires its actin assembly domain for its dot-like localization ^70^. This formin nucleates tropomyosin-decorated linear actin filaments ^71, 72^ required for assembly of the Spitzenkörper. The sole myosin V MyoE, which recruits vesicles to the Spitzenkörper before their delivery at the plasma membrane ^71^, exhibits an almost identical localization to the formin SepA ^72^, whereas vesicles appear to reside somewhat more distally ^73^. These organization similarities suggest that the myosin V may also contribute to focusing the actin structure.

While transport of the actin nucleator by myosin V, as we describe in *S. pombe*, is the simplest scenario to ensure filament barbed end convergence, indirect mechanisms may fulfil the same purpose. In many fungi, the polarisome, a multiprotein complex of still poorly understood stoichiometry, which assembles around the protein Spa2 and regulates formin Bni1 function, plays a key role. In *S. cerevisiae*, where Spa2 phase separates in vitro ^74^, the polarisome component Pea2 binds the myosin V, retaining it at the cell cortex, where motor displacement along actin filaments leads to polarisome spatial focusing ^49^. In turn, polarisome focusing leads to increased formin activity through Spa2-dependent recruitment of Bud6, closing the positive feedback loop ^75^. However, Pea2 is only present in close relatives of *S. cerevisiae* ^50^. In *Neurospora crassa*, the sole myosin V transports Spa2 and its associated <u>c</u>alponin <u>c</u>oiled-coil <u>p</u>rotein-1 (CCP-1) to the Spitzenkörper, thus forming an alternative transport-mediated positive feedback to stabilize F-actin ^76^. In *S. pombe*, we found that Spa2, which lacks the region mediating phase separation in *S. cerevisiae*, does not concentrate at the fusion focus, suggesting it does not play a key role in focus formation. Instead, the *S. pombe* fusion focus offers a simplified model, where components of biomolecular condensation and cytoskeletal feedback converge on the same filament nucleator. This illustrates how evolution, using similar ingredients but tinkering with their use, can produce several solutions using the same organizational principles.

## Materials and methods

### Strain construction

Strains were obtained through standard genetic manipulation of *S. pombe* either by tetrad dissection or transformation and are described in Table S1. All strains are prototroph. Genes were tagged at their endogenous locus, except for Cdc8, Ypt2 and Ypt3, for which an additional copy was expressed by integration of a plasmid at the indicated auxotrophic locus. All plasmids are listed in Table S2. Primers in Table S3.

Gene tagging and deletion at the native genomic locus: Myo52-mScarlet-I:natMX, myo52-mTagBFP2-5xGlyAla-mScarlet-I:natMX, dni1-sfGFP:kanMX tags were produced by PCR-based gene targeting of a fragment from a template pFA6a plasmid containing the appropriate tag and resistance cassette, amplified with primers carrying 5’ extensions corresponding to 78 coding nucleotides at the end of the ORF and 3’UTR, and integrated in the genome by homologous recombination ^77^. Dni2-sfGFP:kanMX and sec6-sfGFP:kanMX, were obtained using plasmids that contained the sfGFP sequence followed by a terminator and resistance marker flanked by the last 500bp from the 3’ end of the ORF without stop codon and 3’UTR of the gene of interest. The Dni2-sfGFP strain lacks the predicted intron. Similarly, myo52Δtail-mScarlet-I:natMX (lacking aa 1183-1516) was obtained using the last 500pb of the truncated sequence. The *fus1^ΔIDR^-sfGFP* allele was transformed in a *fus1* deletion background from a previously available plasmid ^26^. *fus1Δ* and *myo51Δ* strains were similarly constructed using plasmids containing 500bp from the 5’UTR and 3’UTR of the gene of interest flanking a resistance cassette.

Construction of *myo52^AID^*: Myo52-mScarlet-I-3xsAID:natMX was obtained by PCR-based gene targeting of a fragment from a template pFA6a plasmid and crossed to obtain a strain co-expressing Myo52-mScarlet-I-3xsAID:natMX and ura4:padh1:OsTIR1-F74A, as described ^78^. The strain is subsequently referred to as *myo52^AID^*.

Integration of plasmids at exogenous genomic loci: Plasmids for integration at auxotrophic loci were constructed using the single integration vectors described in ^79^. mNeonGreen-cdc8 was introduced as an additional copy under its native promoter at the *leu1+* locus ^40^. GFP-ypt3 and GFP-ypt2 were introduced as additional copies under the *pnmt41* promoter at the *ade6+* and *ura4+* locus respectively. The cytosolic markers (GFP, mCherry and mRaspberry) and LactC2-yeGFP were introduced at the *ura4+* locus under *pact1* promoter. For immunoprecipitation assays, sfGFP, Fus1-sfGFP, Fus1^1–518^–sfGFP and Myo52-3myc were expressed under the *pnmt1* promoter at the *ade6+* locus for GFP-tagged constructs and *lys3+* locus for myc-tagged constructs.

Construction of Fus1^ΔIDR^-FUS^27R^: A plasmid for integration at the *fus1* native locus was constructed by InFusion (pSM3208), containing *fus1* 5’UTR, Fus1 coding sequence for aa 1-491, coding sequence for fused-in-sarcoma aa 1-211 with 27R as described in ^51^, Fus1 coding sequence for aa 792-1372, coding sequence for sfGFP, the kanMX antibiotic resistance cassette and *fus1* 3’ UTR. This was flanked with SalI and SacII restriction sites, used to linearize the fragment and integrate it in the genome by homologous recombination. The Fus1^ΔIDR^-FUS^27R^-K1112A plasmid (pSM3874) was similarly constructed.

### Media and growth condition**s**

Live imaging was adapted from ^80^, with modifications. Cells were pre-cultured in Minimal Sporulation Liquid (MSL) media supplemented with glutamate (MSL+E) as nitrogen source overnight at 30°C. To image mating, *h+* and *h-* cells were mixed at a 1:1 ratio then starved in MSL without nitrogen (MSL-N) for 3-4h at 30°C. 2% electrophoresis-grade agarose pads of MSL+E and MSL-N were used to perform live-cell imaging in vegetative or mating conditions respectively. In short, vegetative cells or 1:1 mixtures of heterothallic *h+* × *h−* cells were placed on top of the pad and covered with a coverslip sealed with VALAP (1:1:1 Vaseline/lanolin/paraffin).

Latrunculin A (LatA; Enzo Life Sciences T-119-0500) was added from a 20mM stock in DMSO to final concentration of 80μM directly in the agarose pad.

For experiments with 5-adamantyl-IAA treatment, liquid media and pads were supplemented with 100nM of 5-adamantyl-IAA (TCI chemicals A3390) prepared from a 1mM stock solution (dissolved in 1 M NaOH and stored at −20°C) and diluted in water. Cells were cultured for 5 hours in liquid media before imaging.

### Microscopy

Imaging to compute the map was performed using a DeltaVision platform (Applied Precision) composed of a customized inverted microscope (IX-71; Olympus), a UPlan Apochromat 100×/1.4 NA oil objective, a camera (CoolSNAP HQ2; Photometrics or 4.2Mpx PrimeBSI sCMOS camera; Photometrics), and a color combined unit illuminator (Insight SSI 7; Social Science Insights). Images were acquired using softWoRx v4.1.2 software (Applied Precision). The Alexa Quad (DAPI/FITC/A594/Cy5) dichroic mirrors filter set was used to record BFP (ex 390/18, em 435/48), RFP (ex 575/25 em 632/60) and GFP (ex 475/20 em 525/50) signals. Cells were imaged every 3 min for 2h30 to 3h, with 100 ms exposure time and 100% intensity for all channels. An additional reference image was taken in DIC (Differential Interference Contrast) for each timepoint using an exposure of 25 ms.

Super resolution imaging in Figure 2B-D was performed on a ZEISS LSM 980 scanning confocal microscope fitted on an inverted Axio Observer 7 Microscope with an Airyscan2 detector optimized for a 63x/1.40 NA oil objective and laser lines 488nm and 561nm. Images were acquired using the Airyscan SR mode (2D SR) and processed with the Zen3.3 (blue edition) software for super resolution. In Figure 2B, C and D, cells were imaged for 1h every 5min using the 561nm laser at 0.4% intensity for RFP signal and 488nm laser at 0.4% for GFP signal, with no averaging. Timepoints represented were taken 5 to 15 min before fusion. For Figure 3 D and S4 C snapshots were taken with same laser lines at 0.8% intensity with an averaging of 2 using the mean intensity method.

Timelapse imaging of vesicles trafficking in Figures 3B and S4A was recorded using a spinning disk microscope composed of an inverted microscope (DMI4000B; Leica) equipped with an HCX Plan Apochromat 100×/1.46 NA oil objective, a CSU22 real-time confocal scanning head (Yokagawa Electric Corporation), solid-state laser lines (Visitron LaserModule 1870), and an electron multiplying charge coupled device camera (iXon-897Life Back Illuminated EMCCD Camera). Time-lapse images were acquired at 260ms or 130ms interval for dual-color or single-color imaging respectively, using the VisiView Premier GOLD Image acquisition Software (Visitron).

Timelapse imaging Fus1^ΔIDR^-FUS^27R^ in Figures 5A-B, 5D-G and movies S4-5, S7-S10 were recorded using a Nikon ECLIPSE Ti2 inverted microscope operated by the NIS Elements software (Nikon). Imaging was performed with a Plan Apochromat Lambda 100X 1.45 NA oil objective and a Prime BSI express sCMOS camera from Teledyne Vision Solutions. Cells were imaged every 600-610 ms for 1 min, with 80 ms exposure time and 10% intensity at 561nm excitation, and 25 ms exposure time and 30% intensity at 488nm excitation with the Lumencor SpectraX light engine (Chroma). An additional reference image was taken in DIC (Differential Interference Contrast) with transmitted light. Timelapse co-imaging of Fus1^ΔIDR^-FUS^LC-27R^ and actin cables labelled with mCherry-LifeAct in Figure 5C and movie S6 were recorded using a spinning disk microscope as stated above.

### Image Analysis for Fusion Focus Mapping and Measurements

All codes for image analysis are available at https://github.com/Valentine-Thomas/FusionFocus-Mapping/tree/master.

Proteins of interest (POI) were included in the following analysis upon two criteria: (i) The proportion of mating pairs fusing during the experiment timeframe was similar to WT and (ii) the POI fluorescence signal was clearly visible after image processing. We also checked that tagging did not grossly affect function by verifying that the length of the fusion process and overall fusion efficiency during imaging was similar to WT.

<u>Method 1:</u> Timelapse movies were processed following the method described in ^58^ to isolate centroid using a custom IJ1 macro (Image_Processing/Centroid-detection_3channels.ijm). Timelapses were split in the three channels and these were aligned using the TurboReg plugin (^81^; https://bigwww.epfl.ch/thevenaz/turboreg/) to limit the influence of stage drift. For each channel, background subtraction was performed using a Rolling Ball algorithm with a radius equal to the width of a cell in pixels. We then corrected for photobleaching, using the Bleach Corrector macro in FIJI (^82^; https://wiki.cmci.info/downloads/bleach_corrector, Image_Processing/corr_bleach.txt). Photobleaching correction was performed by computing the mean fluorescence of all cells in each frame and scaling the fluorescence intensity back to that of the first frame. Images were saved under the name BGN (BackGrouNd subtracted) before cytosolic noise correction. Cytosolic signal was estimated by duplicating the image and applying a median filter with radius equal to the half-width of the POI signal to isolate (measured in pixel, estimated from the size of the signal profile along a line drown across the fusion focus in FIJI and confirmed by assessing spot disappearance by applying the median filter on one timepoint of the timelapse). This filter replaces the signal to isolate by the median value of the background while maintaining cell boundaries. The duplicated image was then subtracted from the original one leading to the obtention of the processed image, saved under the name BGN-MD (MeDian filtered).

In the timelapse, cell fusion events were manually chosen according to two parameters: 1) Myo52 fluorescence signals in the blue and red channel appeared as dots throughout the process, indicating the fusion axis remained in-focus during imaging, and 2) cell-cell fusion occurred during the timelapse. A rectangular Region of Interest (ROI) was defined to encompass the fusion event and used to crop the image for further analysis. An elliptic ROI was defined to measure cytoplasmic fluorescence in one of the partner cells, usually in the partner that did not express the cytosolic signal. Cytosolic signal to determine fusion time was recorded on BGN file before cytosolic signal subtraction using the ‘mCherryIntmeasure’ macro (Image_Processing/mCherryIntmeasure.ijm). Centroid trajectories were tracked on BGN-MD files using the custom ‘i-track’ IJ1 macro (Image_Processing/i_track.ijm), relying on a modified version of ParticleTracker ^58, 83^ using the radius of the POI, a cutoff of 0 and percentile of 0.1. The cutoff allows to ignore high intensity signals while the percentile determines the minimal fluorescence necessary to be detected. We allowed for 3 consecutive missing points in a trajectory (link=3) and a displacement of 15 px between points. As ParticleTracker needs the signal to be (roughly) round, if the POI signal was too wide, only Myo52 references were recorded at this step. For mutants where Myo52 compaction was impacted, see section on compaction mutants Files were formatted for R and Python analysis using the bash code ‘i_track’ (Data_Curation/i_track.sh), which also filtered out all trajectories under 5 timepoints. Manual curation of the trajectories and associated cytosolic fluorescence intensity was performed using the ‘Filter’ script for R for visualization (Data_Curation/Filter.R). In short, fusion events were kept in the analysis pipeline only if (i) trajectories for all channels had been recorded and (ii) fusion time could be accurately determined by our algorithm. The detection of the fusion frame relies on the fitting of a curve using the least squares method. For each timepoint τ, we predicted a curve where y is equal to the mean fluorescence intensity of timepoints 0 to τ for the interval [0, τ] and to the mean fluorescence of timepoints τ + 1 and the last timepoint for the interval [τ + 1, *end*]. The τ for which the error between this prediction and the actual data is minimal is detected as the timepoint preceding the one at which the change in fluorescence occurs and τ + 1 as the first frame after cell fusion. Occasionally, noise from the cell background could lead to the acquisition of multiples trajectories per channel. This noise was removed during the filtering step to keep only one trajectory per channel, recognized by the similarities between channels. The percentage of trajectories kept varied between 50% to 90% depending on the signal intensity.

We computed a correction to reduce the impact of lateral chromatic aberration on our measurements. As centroid coordinates were extracted from cropped ROIs, the coordinates were first transposed to the full field of view using the matlab pipeline ‘coordinate_correction’ (Data_Curation/coordinate_correction.m). Correction for chromatic aberration was obtained using Tetraspeck Microspheres (Invitrogen) as described in ^58^. In short, we measured the shift between the centroid of Tetraspeck beads signals recorded in the green channel, used as reference, and those recorded in blue and red channels, yielding red-to-green and blue-to-green shift matrices over the field of view. These were then applied to the red and blue channel respectively, to correct signal positions using the ‘transform_data’ (Data_Curation/transform_data.m). Bash script ‘SortforPython’ was used to sort trajectories and fluorescent profiles per event (Data_Duration/SortForPython.sh).

Trajectories from all events were aligned in time and in space through the ‘Method1.1_Trajectories-alignment’ jupyter notebook (Method_1/Method1.1_Trajectories-alignment.ipynb). Temporal alignment is performed by detecting the fusion frame and defining it as t=0. Spatial alignment is based on the trajectories of Myo52 in each partner cell. As the signal from *h-* Myo52 is more stable ^19^, we used the center of mass of its trajectory as the origin for a coordinate system. All trajectories were then rotated around the origin such that the ‘fusion axis’, the line passing through both Myo52 trajectories’ center of mass, is on the x-axis. Directed distances along the x-axis (ignoring fluctuations in y, which averaged to 0) and the median fluorescence intensity in the isolated signal were computed using the ‘Method1.2_Computed-Parameters’ notebook (Method_1/Method1.2_Computed-Parameters.ipynb). Plots were obtained using the ‘Method1.3_Plots’ notebook (Method_1/Method1.3_Plots.ipynb).

<u>Method 2:</u> For POI exhibiting a wide signal that could not be analyzed with ‘i-track’, centroid localization was performed using the ‘Method2.1_Spread-signal-coordinates’ jupyter notebook (Method_2/Method2.1_Spread-signal-coordinates.ipynb). In short, the fluorescence profile was extracted from the green channel along a fusion axis computed between the centroid positions of both Myo52 signals acquired through the method described above. Fluorescence profiles were plotted against the x and y coordinates and each fitted with a gaussian curve. When fitting was sufficiently accurate (R>0.6), centroid coordinates were estimated by the center of the gaussian fit. Note that comparison of the two methods, performed for a signal that was round enough to be analyzed by the i-track method, results in overlapping position of the green POI (Figure S1C). Blue and red signal were corrected for chromatic aberration as in Method 1 while green signal was corrected using the ‘Method2.2_POI-signal-correction’ notebook (Method_2/Method2.2_POI-signal-correction.ipynb). The data was then treated similarly to Method 1. This method was used only for detecting plasma membrane localization using LactC2 as seen in Figure S2C.

<u>Method 3:</u> For mutants that heavily impact the compaction of the fusion focus, the wider Myo52 signal could not be detected by the usual ParticleTracker pipeline. We developed another pipeline to map those data. Timelapse images were processed, events selected, and cytoplasmic signal recorded as described above using a variation of the Image processing pipeline (<u>Image_Processing/Centroid-detection-3channels-method3.ijm</u> and itrack_method3.ijm). For each event, a 17 pixel-wide line was manually drawn along the fusion axis and the fluorescence profile along that line was recorded for each timepoint using the ‘d_distance-line-profile’ macro in ImageJ (Method_3/d_distance-line-profile.ijm). Each profile was fitted by a gaussian (R>0.6) to estimate the signal center, and the distance of the center of the fitted curve from the origin of the profile was recorded. These distances were then computed to determine the distance between the different signals and plotted using the ‘Method3_Mutants-mapping’ notebook (Method_3/Method3_Mutants-mapping.ipynb). Note that this largely manual method, used only for *fus1Δ* in Figure 4E, does not correct for chromatic aberrations. However, comparison with the other methods, yielded largely overlapping profiles (Figure S1C).

To measure the lateral distribution of POI at the fusion site (perpendicular to the fusion axis), timelapses were processed as described above. In all cases, we measured lateral distribution in cells crossed to a WT partner. In previous work, we showed that communication between cells influences focus organization ^24^. To ensure we were looking at cell-intrinsic phenotypes, we only took measurements in cells for which the partner WT cell exhibited a round Myo52 signal. At either the last frame before fusion or one of the latest frames at which Myo52 signal in the WT mating partner was still round, a 15 pixel-wide line through the center of Myo52 signal, or POI signal upon *myo52* deletion, was drawn perpendicular to the fusion axis. Fluorescence profiles were extracted using the ‘line-profile’ macro (Spot_Width_Measure/line-profile.ijm). Gaussian fit was performed on the fluorescence profiles (R²>0.6), and the Full Width at Half Maximum (FWHM) of each fitted gaussian used to estimate the width of the different signals. All parameters and plots were obtained using the ‘Spot_width_measure_Gaussian-fitting’ notebook (Spot_Width_Measure/Spot_width_measure_Gaussian-fitting.ipynb).

To estimate the precision of distance measurements, we considered two main sources of error. First, chromatic aberrations, though partially corrected by microscope lenses, are a source of error for fluorophores at distinct emission wavelength. By using tagged Myo52 emitting in two distinct wavelengths, we measured a residual distance after correction for chromatic aberration of 4nm between GFP and RFP (Figure 1E), which directly affects our POI-GFP to Myo52-mScarlet distance measurements. We also measured a 10 nm residual distance between GFP and BFP channels (Figure 1F), which does not directly affect distance measurements, but impacts the precise positioning of the fusion axis, both in x-y and in z. Indeed, if the axis, along which we measure distances, is placed at an angle with where the real fusion axis should be, this will also affect the precision of the measurements. In the x-y plane, the maximal imprecision of axis position occurs if Myo52-BFP real position is in fact displaced in y. For a displacement in y of 10 nm and an average distance between Myo52 foci of 350 nm, this results in a negligeable 0.04% underestimation. The error in z is more difficult to estimate. Grossly out-of-focus signals were omitted from analysis by restricting it to events where both Myo52 appeared as dots throughout the process, but slight displacement in z in those kept for analysis will also skew the position of the fusion axis. From z stack imaging, we estimate that the maximal z distance between the two Myo52 foci in events kept for analysis was 100 nm, which, for an average distance between Myo52 foci of 350 nm, would result in a 4% underestimation of the real distance along the fusion axis. Thus, for distances between red and green signals, we estimate the maximal error to 4% due to underestimation due to axis placement, to be added to 4nm due to chromatic aberration. For a measured distance of 100 nm (all distances are shorter than this value, except for Cdc8 and Myo51), the total maximal error (chromatic aberrations and axis positioning) is about 8 nm.

### Statistics

For each POI, to determine whether its position was changing relative to the Myo52 reference over time, we used linear regression on data from timepoint [-60,0] to determine if the slope was significantly different from 0. To determine if the POI’s localization was significantly different from Myo52, we tested whether the mean distance between the POI and Myo52 at early and late timepoints ([-60,-30] or [-27,0] pooled together, respectively) was significantly different from 0 using the Wilcoxon signed-rank test. Statistical results, as well as the number of trajectories quantified for each protein, are displayed in Table S4.

To compare protein width, we performed a Mann-Whitney U test on each relevant pair of groups.

To test for differences in Fus1^ΔIDR^-FUS^LC-27R^-sfGFP long-range track frequency, we used a paired t-test.

### Co-immunoprecipitation

Cells were first pre-cultured overnight in 50 mL of MSL+ammonium media (MSL+N) supplemented with 5ng/mL of thiamine at 30°C, then washed and diluted to around OD600 = 0.07 into MSL+N and incubated for 7-8 hours at 30°C then diluted once more to OD600 = 0.05 and incubated overnight at 30°C until OD600 = 0.6 to 0.8. Then the cells were washed with cold lysis buffer (50mM Tris-HCl pH 7.4, 200mM NaCl, 1mM EDTA, with protease inhibitors (antipain 0.7µM, aprotinin 0.08 µM, chymostatin C 0.8 µM, leupeptin 1µM, pepstatin A 0.7 µM, 1.10 phenanthroline 0.5 mM, benzamidine-HCl 3µM, PMSF 100µM; final concentration indicated) and resuspended in lysis buffer supplemented with detergent (50mM Tris-HCl pH 7.4, 200mM NaCl, 1mM EDTA, 1% LMNG, 0.1% CHS with protease inhibitors). Cells were then lysed using a MagNA Lyser (Roche), using 4 cycles of 6500 rpm for 45 sec. Lysis efficiency was checked by light microscopy, if efficiency was inferior to 60% of the cells, an extra round of MagNA lyser was performed. Supernatant was then collected by piercing the tubes, placed inside another tube, and centrifugation at 13’000 rpm for 20 min (4°C). Total protein amount was equalized (∼10’000ng) and added to 20 μl of GFP-Trap magnetic beads (Roche) pre-washed in lysis buffer (50mM Tris-HCl pH 7.4, 200mM NaCl, 1mM EDTA, 1% Tergitol with protease inhibitors). Proteins were IP for 3h before being washed three times in lysis buffer. Input and IP samples were diluted in 4x and 1.5X NuPAGE LDS sample buffer (Invitrogen) with β-mercaptoethanol (Sigma), respectively, and boiled for 15 min at 95 °C. Samples were run on 8% SDS-PAGE (GenScript), blotted, and detected with either anti-GFP antibody from mouse (Roche 11814460001) and HRP conjugated anti-mouse secondary antibody (Promega W402B) or anti-Myc antibody from goat (Abcam ab9132) and donkey anti-goat secondary antibody (Santa Cruz sc-2354).

## Acknowledgements

We thank Dr Brieuc Chauvin for the first observation of Fus1^ΔIDR^-FUS^LC-27R^-sfGFP mitotic foci and Wanlan Li for help with Python. This work was supported by an ERC Advanced Grant (SexYeast) and Swiss National Science Foundation grants #176396 and #191990 to SGM and #212288 to MK. HM was supported by a Swiss Government Excellence Scholarship.

## Author contributions

The project was designed by SGM and VT. VT developed the centroid-tracking method with help of AP and MK and did all experiments except those in Figure 5. HM performed the experiments in Figure 5. LM provided technical help for cloning, strain generation and immunoprecipitations. SGM and MK acquired funding. SGM managed the project, provided supervision and wrote the manuscript, which was reviewed by all authors.

## Figure legends

**Figure S1:**
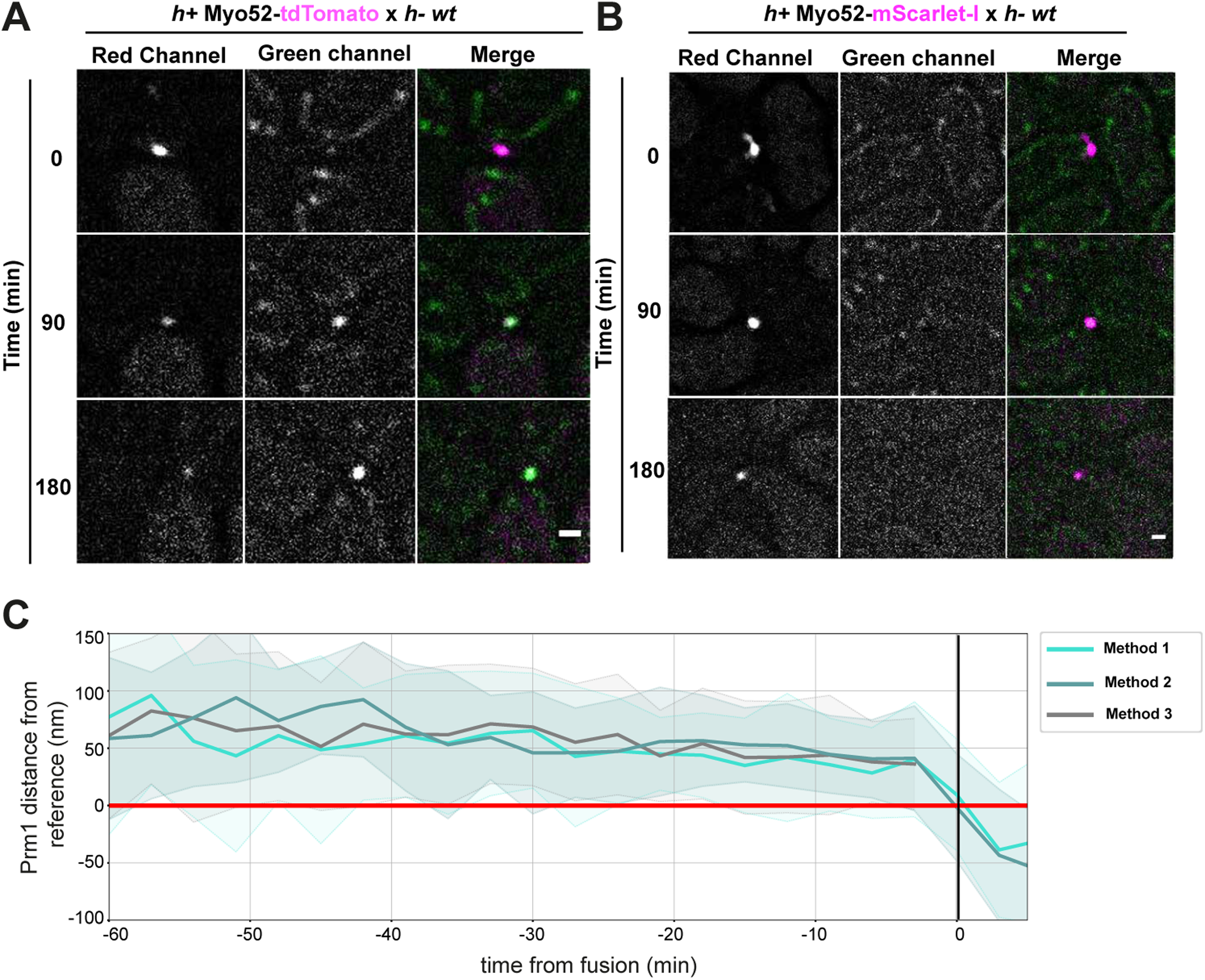
tdTomato photoconversion and comparison of measurement methods A-B. RFP, GFP channel and merge timelapse images of yeast mating cells expressing only Myo52-tdTomato (left) or only Myo52-mScarlet-I (right). The cells were imaged every 3 min. Note detection of tdTomato but not mScarlet-I over time in the GFP channel. **C.** Median distances of Prm1 from Myo52 measured with either method 1 (centroid tracking of Myo52 references and Prm1) used for all round signals, method 2 (centroid tracking of only Myo52 references) used for wide signals or method 3 (manual fusion axis placement) used in mutants where Myo52 reference was not round. Shaded zones show standard deviation, black line shows fusion time. Scale bars are 1 µm.

**Figure S2:**
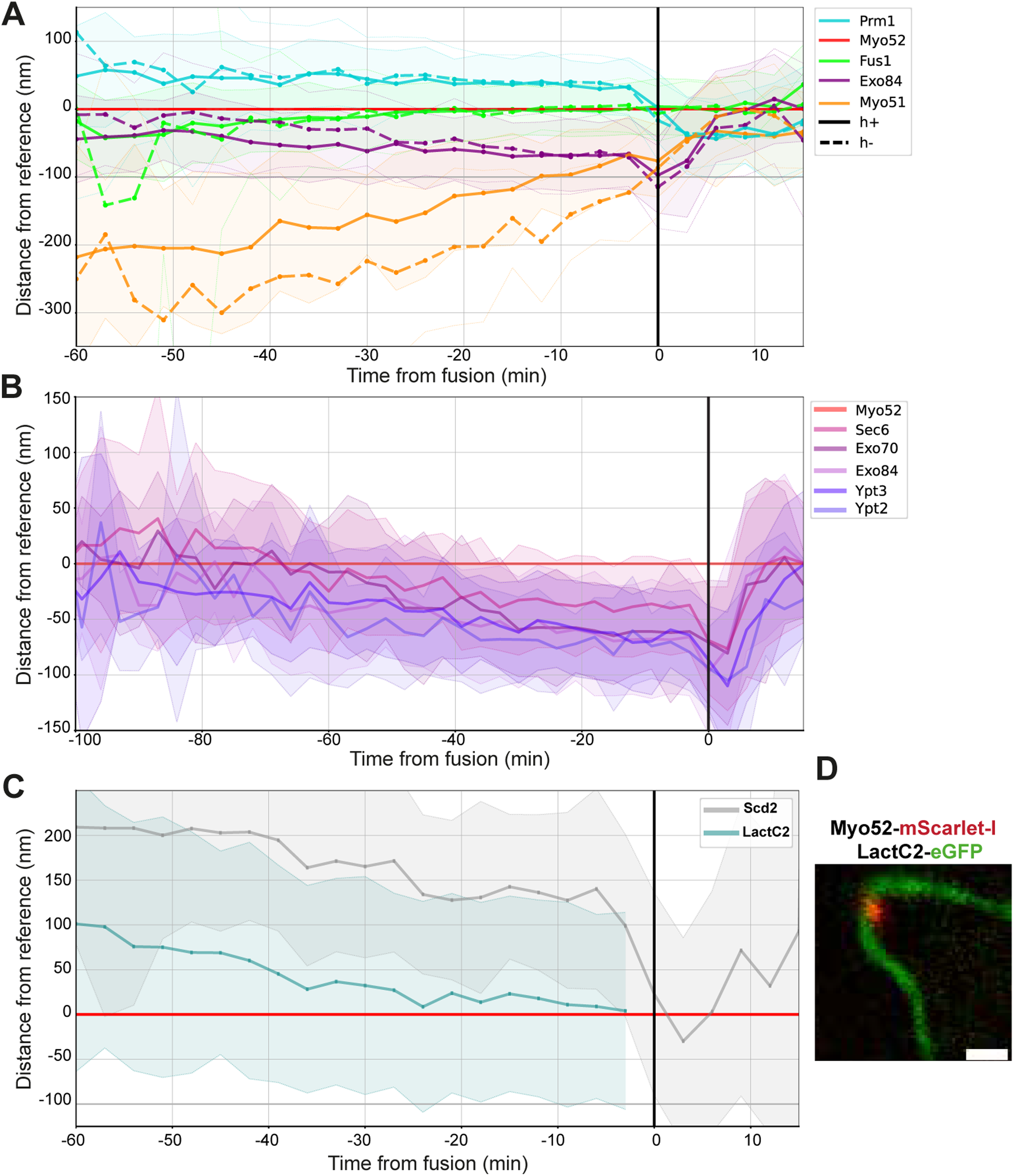
Comparison of *h+* and *h-* cells and LactC2 localization. **A.** Comparison of median directed distances between selected proteins and Myo52 in *h+* (plain lines) and *h-* (dashed lines) cells over time. **B.** Median directed distances from Myo52 of exocyst subunits and Rab GTPases labelling secretory vesicles over time, as in Figure 2D but from-100 min before fusion time. **C.** Median directed distance of LactC2 from Myo52 over time. The Scd2 profile is shown for comparison. **D.** Deltavision image of LactC2-GFP and Myo52-mScarlet-I distribution during fusion. On all graphs, shaded zones show standard deviation, black line shows fusion time. Scale bar is 1 µm.

**Figure S3:**
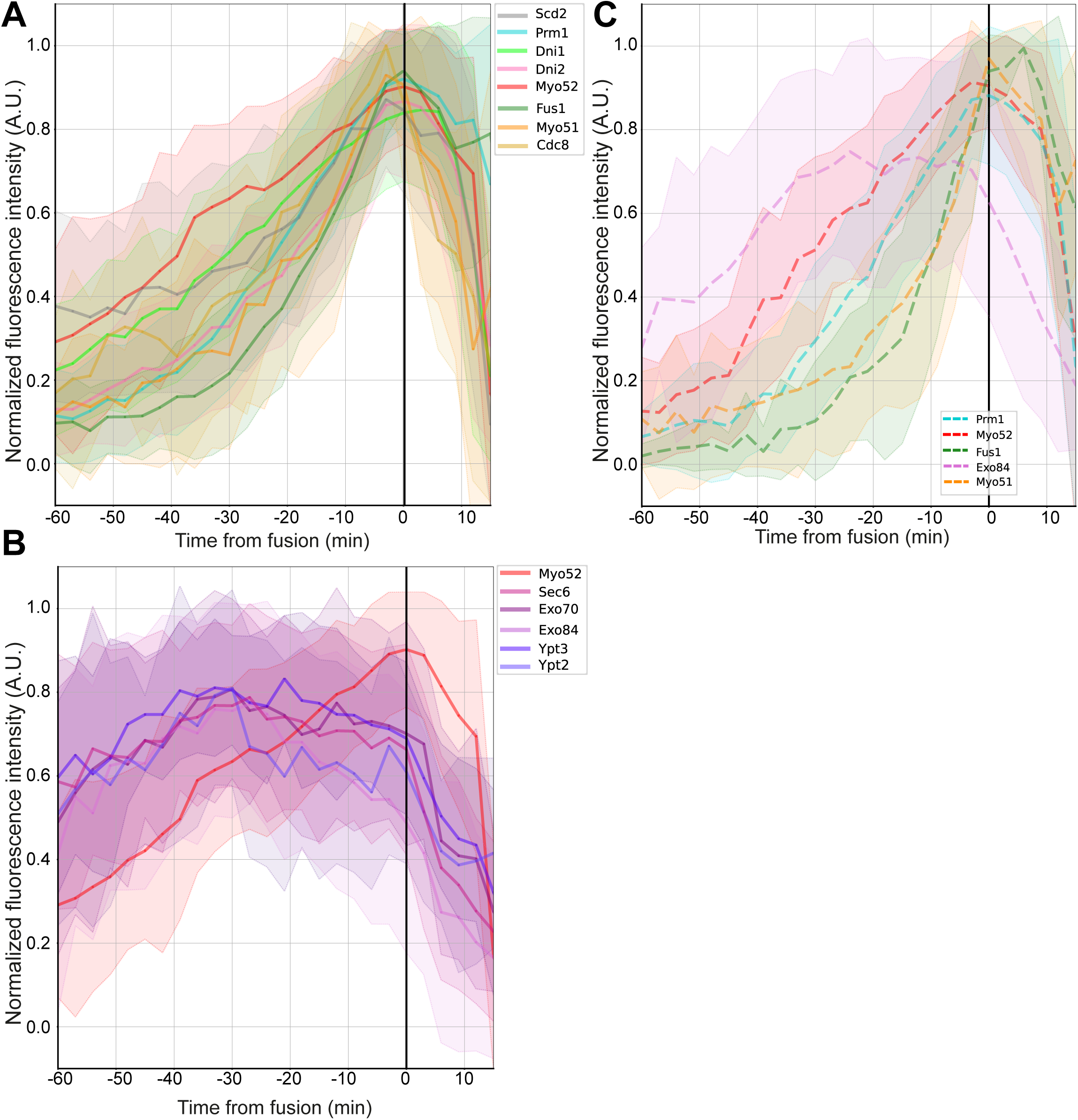
Evolution of fluorescence intensity at the fusion site over time A-B. Fluorescence intensity of indicated proteins at the fusion focus in *h+* cells during the fusion process in proteins showing gradual increase until fusion time (A) and in secretory vesicle-associated factors (B). The Myo52 fluorescence profile is shown on both graphs for reference. **C.** Fluorescence intensity of selected proteins at the fusion focus in *h-* cells. On all graphs, shaded zones show standard deviation, black line shows fusion time.

**Figure S4:**
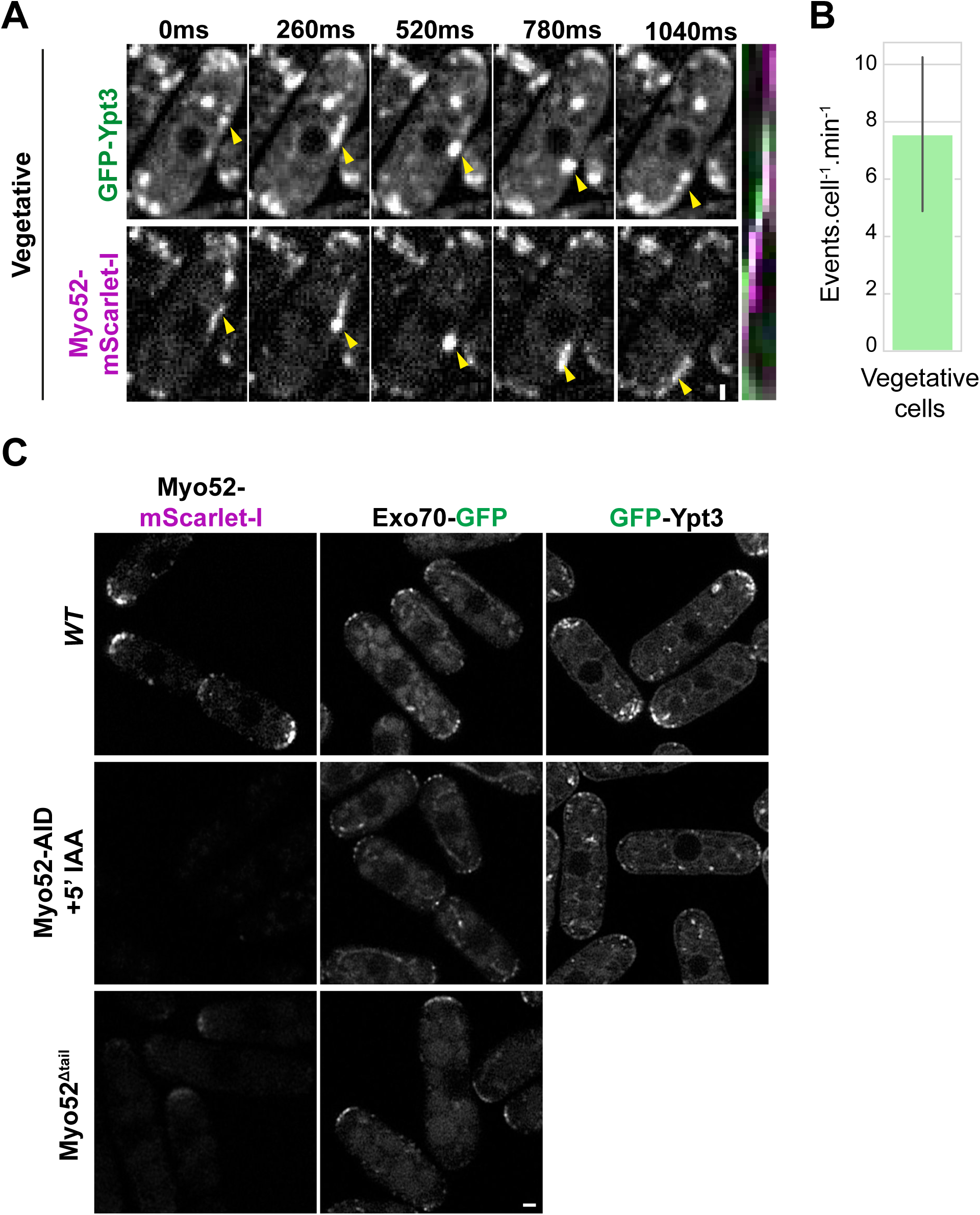
Ypt3 and Exo70 localization in vegetative WT and *myo52* mutant cells. **A.** Spinning disk two-color timelapse images of Ypt3 and Myo52 linear movements in proliferating mitotic cells. Yellow arrowheads track an individual vesicle. A kymograph is shown on the right. The graph shows the average number of linear movements observed per cell per minute over at least 4 consecutive time points. **B.** Airyscan images of Myo52-mScarlet-I, Exo70-GFP and GFP-Ypt3 in indicated mitotically proliferating cells. Myo52^AID^ and Myo52^Δtail^ are also tagged with mScarlet-I. Note disappearance of Myo52^AID^. Scale bars are 1 µm.

**Figure S5:**
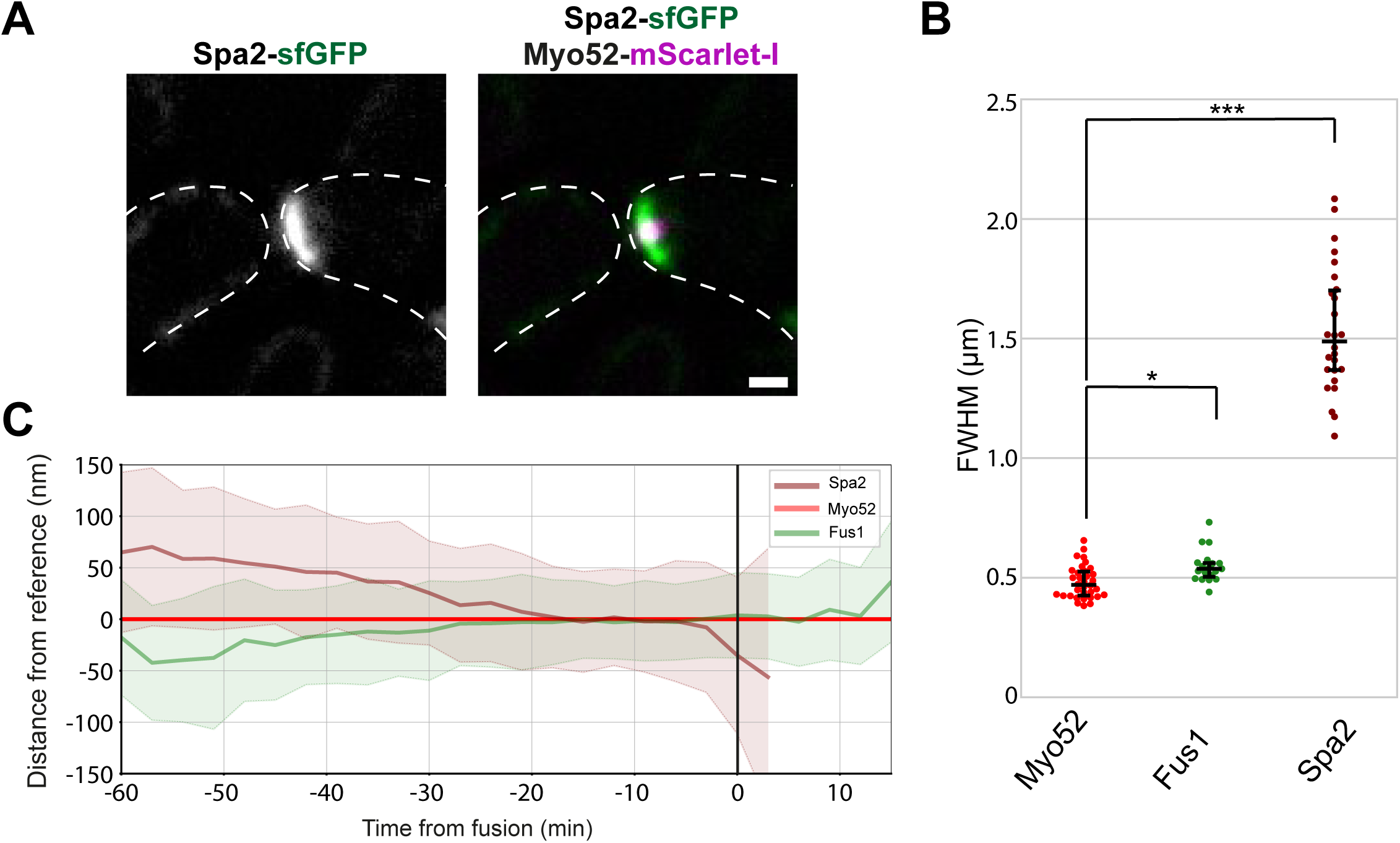
Spa2 distribution at the fusion site. **A.** Spa2-sfGFP distribution in WT *h+* cells also expressing Myo52-mScarlet-I mated with WT *h-* cells. **B.** FWHM of Myo52-mScarlet-I, Fus1-sfGFP and Spa2-sfGFP fluorescence profiles perpendicular to fusion axis in cells at the last time point before fusion. The data for Fus1 is shown for comparison and is identical to that of Figure 6B. **C.** Median directed distances from Myo52 of Fus1 and Spa2 over the fusion process computed with method 1 and 2, respectively. The data for Fus1 is show for comparison and is identical to that of Figure 2A-B. Scale bar is 1 µm.

**Figure S6:**
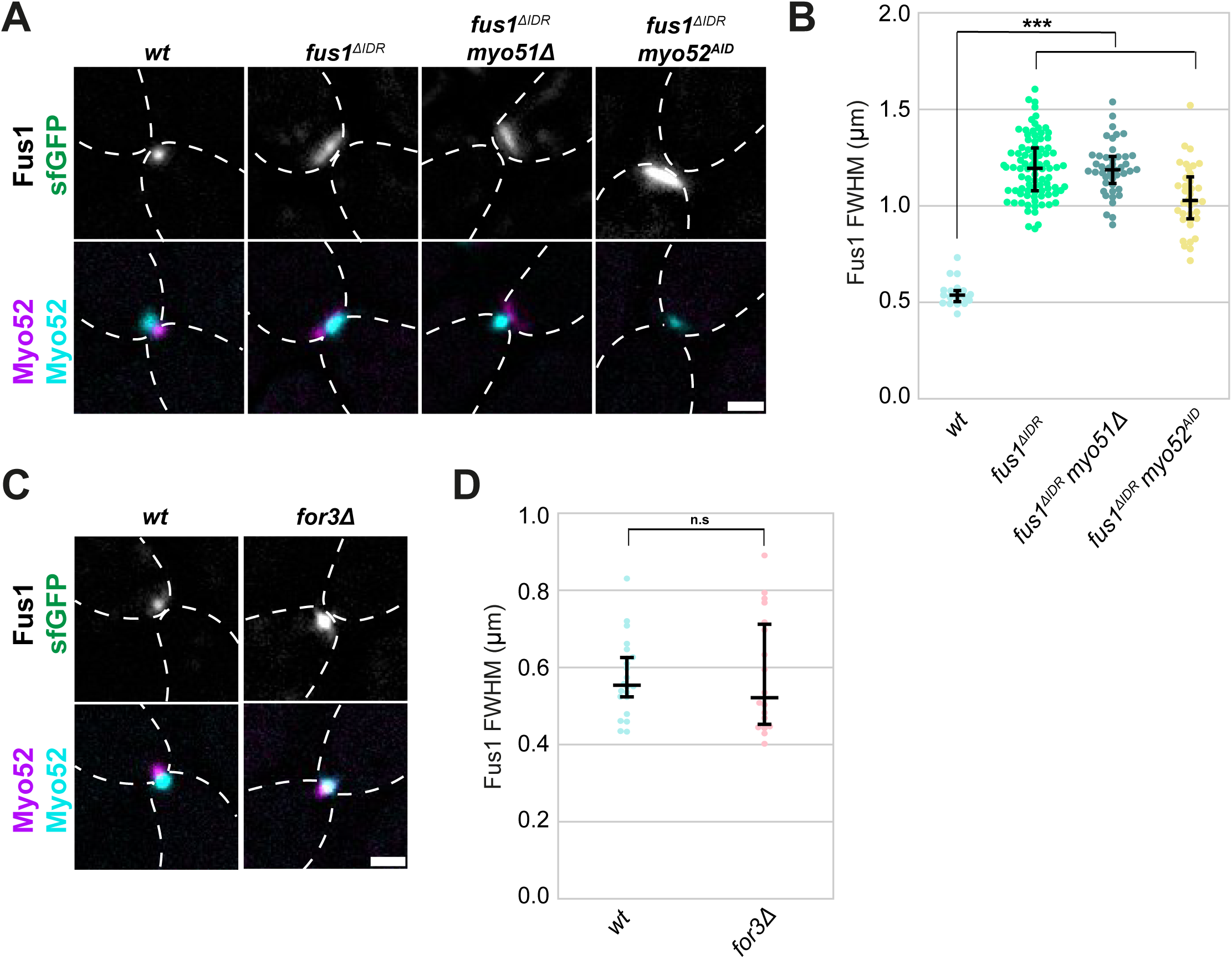
Fus1 distribution in *fus1^ΔIDR^* and *for3Δ*. **A.** Fus1-sfGFP (left) and Fus1^ΔIDR^-sfGFP distribution in *h+* cells that are either WT, *myo52^AID^* or *myo51Δ*. The bottom row shows the Myo52-mScarlet-I signal in the same cell and the Myo52-mTagBFP2 signal in the partner WT cell. Note absence of Myo52^AID^-mScarlet-I signal. **B.** FWHM of Fus1-sfGFP fluorescence profiles perpendicular to fusion axis in cells as in (A) at the last time point before fusion or the last time point when the Myo52-mTagBFP2 signal was round. **C.** Fus1-sfGFP distribution in *h-* cells that are either WT or *for3Δ*. The bottom row shows the Myo52-mScarlet-I signal in the same cell and the Myo52-mTagBFP2 signal in the partner WT cell. **D.** FWHM of Fus1-sfGFP fluorescence profiles perpendicular to fusion axis in cells as in (C) at the last time point before fusion. Scale bars are 1 µm.

---

**Movie S1: Timelapse of GFP-Ypt3 in early mating**

Images were taken at 130 ms interval and are displayed in real time. Time is indicated from start of imaging.

**Movie S2: Timelapse of GFP-Ypt3 during the fusion process**

Images were taken at 130 ms interval and are displayed in real time. Time is indicated from start of imaging.

**Movie S3: Timelapse of GFP-Ypt3 in myo52^AID^ cells during the fusion process**

The timelapse shows *h+ myo52^AID^-mScarlet-I* cells expressing GFP-Ypt3 crossed to *h-* WT cells expressing cytosolic mRaspberry. The cells were treated with 100 nM 5-adamantyl-IAA (5’IAA) during starvation for 5h. Note that GFP-Ypt3 fails to localize to the fusion focus before cell fusion but is recruited by Myo52 expressed in the WT cells (from 27 min) after pore opening visible by the transfer of mRaspberry to the *h+* cell (starting at 21 min). Time is in min from start of imaging. Scale bar is 1µm.

**Movie S4: Long-range linear movement of Fus1^ΔIDR^-FUS^LC-27R^ with Myo52**

Timelapse images of Fus1^ΔIDR^-FUS^LC-27R^-sfGFP and Myo52-mScarlet-I in vegetative cells. This corresponds to the cell shown in Figure 5A. Images were taken at 600 ms interval and are sped up 7.2-fold. Scale bar is 2µm.

**Movie S5: Further examples of long-range linear movements of Fus1^ΔIDR^-FUS^LC-27R^ with Myo52**

Timelapse images of Fus1^ΔIDR^-FUS^LC-27R^-sfGFP and Myo52-mScarlet-I in vegetative cells. Images were taken at 600 ms interval and are sped up 7.2-fold. The shorter time lapses stop on the last frame. Scale bar is 2µm.

**Movie S6: Movement of Fus1^ΔIDR^-FUS^LC-27R^ on an actin cable**

Spinning disk timelapse images of Fus1^ΔIDR^-FUS^LC-27R^-sfGFP and LifeAct-mCherry in vegetative cells. The arrowhead tracks a Fus1 dot. Images were taken at 500 ms interval and are sped up 6-fold. Scale bar is 2µm.

**Movie S7: Linear movement of Myo52 with and beyond Fus1^ΔIDR^-FUS^LC-27R^**

Timelapse images of Fus1^ΔIDR^-FUS^LC-27R^-sfGFP and Myo52-mScarlet-I in vegetative cells. This corresponds to the cell shown in Figure 5D. Images were taken at 600 ms interval and are sped up 7.2-fold. Yellow and cyan arrowheads point to two Myo52 dots apparently emanating from the Fus1^ΔIDR^-FUS^LC-27R^ focus. Scale bar is 2µm.

**Movie S8: Further examples of linear movements of Myo52 with and beyond Fus1^ΔIDR^-_FUS_LC-27R.**

Timelapse images of Fus1^ΔIDR^-FUS^LC-27R^-sfGFP and Myo52-mScarlet-I in vegetative cells. Note the Myo52 dots apparently continuing on a linear track beyond the Fus1^ΔIDR^-FUS^LC-27R^ focus. The yellow arrowheads point to continuing Myo52 dots and semi-static Fus1 dots in the respective channels. Images were taken at 600 ms interval and are sped up 7.2-fold. Scale bar is 2µm.

**Movie S9: Convergent movement of Myo52-decorated Fus1^ΔIDR^-FUS^LC-27R^ foci.**

Timelapse images of Fus1^ΔIDR^-FUS^LC-27R^-sfGFP and Myo52-mScarlet-I in vegetative cells. This corresponds to the cell shown in Figure 5E. Images were taken at 600 ms interval and are sped up 7.2-fold. Scale bar is 2µm.

**Movie S10: Further examples of convergent movements of Myo52-decorated Fus1^ΔIDR^-FUS^LC-27R^ foci.**

Timelapse images of Fus1^ΔIDR^-FUS^LC-27R^-sfGFP and Myo52-mScarlet-I in vegetative cells. Images were taken at 600 ms interval and are sped up 7.2-fold. The shorter time lapses stop on the last frame. Scale bar is 2µm.

**Table S1: Strains used in this study**

Strains are indicated in the first Figure where they appear. The methods column indicates how they were obtained. Please refer to Tables S2 and S3 for details of the plasmids and primers.

**Table S2: Plasmids used in this study**

**Table S3: Primers used in this study**

**Table S4: Statistics and number of profiles per localized protein**

The table shows the number of profiles measured for each tagged protein, organized by Figure presentation. For the profiles in Figure 2, linear regression statistics comparing the slopes of the curves of the whole hour pre-fusion [-60,0], or each half-hour [-60,-30] and [-27,0] to that of the Myo52 reference are provided. Note that Fus1 and Myo52 traces are not significantly different in the last half-hour.

## References

1. Sieber, B., Coronas-Serna, J.M. & Martin, S.G. A focus on yeast mating: From pheromone signaling to cell-cell fusion. Semin Cell Dev Biol 133, 83–95 (2023).

2. Gingras, R.M., Sulpizio, A.M., Park, J. & Bretscher, A. High-resolution secretory timeline from vesicle formation at the Golgi to fusion at the plasma membrane in S. cerevisiae. eLife 11 (2022).

3. Casavola, E.C. et al. Ypt32p and Mlc1p bind within the vesicle binding region of the class V myosin Myo2p globular tail domain. Mol Microbiol 67, 1051–1066 (2008).

4. Lipatova, Z. et al. Direct interaction between a myosin V motor and the Rab GTPases Ypt31/32 is required for polarized secretion. Mol Biol Cell 19, 4177–4187 (2008).

5. Santiago-Tirado, F.H., Legesse-Miller, A., Schott, D. & Bretscher, A. PI4P and Rab inputs collaborate in myosin-V-dependent transport of secretory compartments in yeast. Dev Cell 20, 47–59 (2011).

6. Jin, Y. et al. Myosin V transports secretory vesicles via a Rab GTPase cascade and interaction with the exocyst complex. Dev Cell 21, 1156–1170 (2011).

7. Lo Presti, L. & Martin, S.G. Shaping fission yeast cells by rerouting actin-based transport on microtubules. Curr Biol 21, 2064–2069 (2011).

8. Schott, D.H., Collins, R.N. & Bretscher, A. Secretory vesicle transport velocity in living cells depends on the myosin-V lever arm length. J Cell Biol 156, 35–39 (2002).

9. Motegi, F., Arai, R. & Mabuchi, I. Identification of two type V myosins in fission yeast, one of which functions in polarized cell growth and moves rapidly in the cell. Mol Biol Cell 12, 1367–1380 (2001).

10. Feierbach, B. & Chang, F. Roles of the fission yeast formin for3p in cell polarity, actin cable formation and symmetric cell division. Curr Biol 11, 1656–1665 (2001).

11. Bendezu, F.O., Vincenzetti, V. & Martin, S.G. Fission yeast Sec3 and Exo70 are transported on actin cables and localize the exocyst complex to cell poles. PLoS One 7, e40248 (2012).

12. Donovan, K.W. & Bretscher, A. Myosin-v is activated by binding secretory cargo and released in coordination with rab/exocyst function. Dev Cell 23, 769–781 (2012).

13. Donovan, K.W. & Bretscher, A. Tracking individual secretory vesicles during exocytosis reveals an ordered and regulated process. J Cell Biol 210, 181–189 (2015).

14. Clayton, J.E. et al. Fission yeast tropomyosin specifies directed transport of myosin-V along actin cables. Mol Biol Cell 25, 66–75 (2014).

15. Hodges, A.R. et al. Tropomyosin is essential for processive movement of a class V myosin from budding yeast. Curr Biol 22, 1410–1416 (2012).

16. Schuh, M. An actin-dependent mechanism for long-range vesicle transport. Nat Cell Biol 13, 1431–1436 (2011).

17. Martin, S.G. & Arkowitz, R.A. Cell polarization in budding and fission yeasts. FEMS microbiology reviews 38, 228–253 (2014).

18. Lo Presti, L., Chang, F. & Martin, S.G. Myosin Vs organize actin cables in fission yeast. Mol Biol Cell 23, 4579–4591 (2012).

19. Dudin, O. et al. A formin-nucleated actin aster concentrates cell wall hydrolases for cell fusion in fission yeast. J Cell Biol 208, 897–911 (2015).

20. Petersen, J., Weilguny, D., Egel, R. & Nielsen, O. Characterization of fus1 of Schizosaccharomyces pombe: a developmentally controlled function needed for conjugation. Mol Cell Biol 15, 3697–3707 (1995).

21. Billault-Chaumartin, I. et al. Actin assembly requirements of the formin Fus1 to build the fusion focus. J Cell Sci 135 (2022).

22. Win, T.Z., Gachet, Y., Mulvihill, D.P., May, K.M. & Hyams, J.S. Two type V myosins with non-overlapping functions in the fission yeast Schizosaccharomyces pombe: Myo52 is concerned with growth polarity and cytokinesis, Myo51 is a component of the cytokinetic actin ring. J Cell Sci 114, 69–79 (2001).

23. Muriel, O., Michon, L., Kukulski, W. & Martin, S.G. Ultrastructural plasma membrane asymmetries in tension and curvature promote yeast cell fusion. bioRxiv, 2021.2003.2018.435973 (2021).

24. Dudin, O., Merlini, L. & Martin, S.G. Spatial focalization of pheromone/MAPK signaling triggers commitment to cell-cell fusion. Genes Dev 30, 2226–2239 (2016).

25. Merlini, L. et al. Inhibition of Ras activity coordinates cell fusion with cell-cell contact during yeast mating. J Cell Biol 217, 1467–1483 (2018).

26. Billault-Chaumartin, I., Muriel, O., Michon, L. & Martin, S.G. Condensation of the fusion focus by the intrinsically disordered region of the formin Fus1 is essential for cell-cell fusion. Curr Biol 32, 4752–4761 e4710 (2022).

27. Tang, Q. et al. A single-headed fission yeast myosin V transports actin in a tropomyosin-dependent manner. J Cell Biol 214, 167–179 (2016).

28. Dudin, O. et al. A systematic screen for morphological abnormalities during fission yeast sexual reproduction identifies a mechanism of actin aster formation for cell fusion. PLoS genetics 13, e1006721 (2017).

29. Curto, M.A. et al. Membrane Organization and Cell Fusion During Mating in Fission Yeast Requires Multi-Pass Membrane Protein Prm1. Genetics (2014).

30. Billault-Chaumartin, I. & Martin, S.G. Capping Protein Insulates Arp2/3-Assembled Actin Patches from Formins. Curr Biol 29, 3165–3176 e3166 (2019).

31. Vjestica, A., Merlini, L., Nkosi, P.J. & Martin, S.G. Gamete fusion triggers bipartite transcription factor assembly to block re-fertilization. Nature 560, 397–400 (2018).

32. Kurahashi, H., Imai, Y. & Yamamoto, M. Tropomyosin is required for the cell fusion process during conjugation in fission yeast. Genes Cells 7, 375–384 (2002).

33. Clemente-Ramos, J.A. et al. The tetraspan protein Dni1p is required for correct membrane organization and cell wall remodelling during mating in Schizosaccharomyces pombe. Mol Microbiol 73, 695–709 (2009).

34. Curto, M.A., Moro, S., Yanguas, F., Gutierrez-Gonzalez, C. & Valdivieso, M.H. The ancient claudin Dni2 facilitates yeast cell fusion by compartmentalizing Dni1 into a membrane subdomain. Cellular and molecular life sciences: CMLS 75, 1687–1706 (2018).

35. Doyle, A., Martin-Garcia, R., Coulton, A.T., Bagley, S. & Mulvihill, D.P. Fission yeast Myo51 is a meiotic spindle pole body component with discrete roles during cell fusion and spore formation. J Cell Sci 122, 4330–4340 (2009).

36. Picco, A., Mund, M., Ries, J., Nedelec, F. & Kaksonen, M. Visualizing the functional architecture of the endocytic machinery. eLife 4 (2015).

37. Wang, L., Jackson, W.C., Steinbach, P.A. & Tsien, R.Y. Evolution of new nonantibody proteins via iterative somatic hypermutation. Proc Natl Acad Sci U S A 101, 16745–16749 (2004).

38. Chang, E.C. et al. Cooperative interaction of S. pombe proteins required for mating and morphogenesis. Cell 79, 131–141 (1994).

39. Bendezu, F.O. & Martin, S.G. Cdc42 Explores the Cell Periphery for Mate Selection in Fission Yeast. Curr Biol 23, 42–47 (2013).

40. Hatano, T. et al. mNG-tagged fusion proteins and nanobodies to visualize tropomyosins in yeast and mammalian cells. J Cell Sci 135 (2022).

41. Wheatley, E. & Rittinger, K. Interactions between Cdc42 and the scaffold protein Scd2: requirement of SH3 domains for GTPase binding. Biochem J 388, 177–184 (2005).

42. Haupt, A. & Minc, N. Gradients of phosphatidylserine contribute to plasma membrane charge localization and cell polarity in fission yeast. Mol Biol Cell 28, 210–220 (2017).

43. Fairn, G.D., Hermansson, M., Somerharju, P. & Grinstein, S. Phosphatidylserine is polarized and required for proper Cdc42 localization and for development of cell polarity. Nat Cell Biol 13, 1424–1430 (2011).

44. Cheng, H. et al. Role of the Rab GTP-binding protein Ypt3 in the fission yeast exocytic pathway and its connection to calcineurin function. Mol Biol Cell 13, 2963–2976 (2002).

45. Craighead, M.W., Bowden, S., Watson, R. & Armstrong, J. Function of the ypt2 gene in the exocytic pathway of Schizosaccharomyces pombe. Mol Biol Cell 4, 1069–1076 (1993).

46. Merlini, L. et al. Local Pheromone Release from Dynamic Polarity Sites Underlies Cell-Cell Pairing during Yeast Mating. Curr Biol 26, 1117–1125 (2016).

47. Bendezu, F.O. & Martin, S.G. Actin cables and the exocyst form two independent morphogenesis pathways in the fission yeast. Mol Biol Cell 22, 44–53 (2011).

48. Mulvihill, D.P., Edwards, S.R. & Hyams, J.S. A critical role for the type V myosin, Myo52, in septum deposition and cell fission during cytokinesis in Schizosaccharomyces pombe. Cell Motil Cytoskeleton 63, 149–161 (2006).

49. Dunkler, A. et al. Type V myosin focuses the polarisome and shapes the tip of yeast cells. J Cell Biol 220 (2021).

50. Lichius, A., Yanez-Gutierrez, M.E., Read, N.D. & Castro-Longoria, E. Comparative live-cell imaging analyses of SPA-2, BUD-6 and BNI-1 in Neurospora crassa reveal novel features of the filamentous fungal polarisome. PLoS One 7, e30372 (2012).

51. Wang, J. et al. A Molecular Grammar Governing the Driving Forces for Phase Separation of Prion-like RNA Binding Proteins. Cell 174, 688–699 e616 (2018).

52. Sakamoto, S. et al. mDia1/3 generate cortical F-actin meshwork in Sertoli cells that is continuous with contractile F-actin bundles and indispensable for spermatogenesis and male fertility. PLoS Biol 16, e2004874 (2018).

53. Scott, B.J., Neidt, E.M. & Kovar, D.R. The functionally distinct fission yeast formins have specific actin-assembly properties. Mol Biol Cell 22, 3826–3839 (2011).

54. Boyd, C., Hughes, T., Pypaert, M. & Novick, P. Vesicles carry most exocyst subunits to exocytic sites marked by the remaining two subunits, Sec3p and Exo70p. J Cell Biol 167, 889–901 (2004).

55. Finger, F.P., Hughes, T.E. & Novick, P. Sec3p is a spatial landmark for polarized secretion in budding yeast. Cell 92, 559–571 (1998).

56. Roumanie, O. et al. Rho GTPase regulation of exocytosis in yeast is independent of GTP hydrolysis and polarization of the exocyst complex. J Cell Biol 170, 583–594 (2005).

57. Heider, M.R. et al. Subunit connectivity, assembly determinants and architecture of the yeast exocyst complex. Nature structural & molecular biology 23, 59–66 (2016).

58. Picco, A. et al. The In Vivo Architecture of the Exocyst Provides Structural Basis for Exocytosis. Cell 168, 400–412 e418 (2017).

59. Ahmed, S.M. et al. Exocyst dynamics during vesicle tethering and fusion. Nat Commun 9, 5140 (2018).

60. Riquelme, M. et al. The Neurospora crassa exocyst complex tethers Spitzenkorper vesicles to the apical plasma membrane during polarized growth. Mol Biol Cell 25, 1312–1326 (2014).

61. Sivaram, M.V., Saporita, J.A., Furgason, M.L., Boettcher, A.J. & Munson, M. Dimerization of the exocyst protein Sec6p and its interaction with the t-SNARE Sec9p. Biochemistry 44, 6302–6311 (2005).

62. Jourdain, I., Dooley, H.C. & Toda, T. Fission yeast sec3 bridges the exocyst complex to the actin cytoskeleton. TraYic 13, 1481–1495 (2012).

63. Borgal, L. & Wakefield, J.G. Context-dependent spindle pole focusing. Essays Biochem 62, 803–813 (2018).

64. Raff, J.W. Phase Separation and the Centrosome: A Fait Accompli? Trends Cell Biol 29, 612–622 (2019).

65. Heald, R., Tournebize, R., Habermann, A., Karsenti, E. & Hyman, A. Spindle assembly in Xenopus egg extracts: respective roles of centrosomes and microtubule self-organization. J Cell Biol 138, 615–628 (1997).

66. Gaglio, T., Dionne, M.A. & Compton, D.A. Mitotic spindle poles are organized by structural and motor proteins in addition to centrosomes. J Cell Biol 138, 1055–1066 (1997).

67. Merdes, A., Ramyar, K., Vechio, J.D. & Cleveland, D.W. A complex of NuMA and cytoplasmic dynein is essential for mitotic spindle assembly. Cell 87, 447–458 (1996).

68. Hueschen, C.L., Kenny, S.J., Xu, K. & Dumont, S. NuMA recruits dynein activity to microtubule minus-ends at mitosis. eLife 6 (2017).

69. Riquelme, M. & Sanchez-Leon, E. The Spitzenkorper: a choreographer of fungal growth and morphogenesis. Curr Opin Microbiol 20, 27–33 (2014).

70. Sharpless, K.E. & Harris, S.D. Functional characterization and localization of the Aspergillus nidulans formin SEPA. Mol Biol Cell 13, 469–479 (2002).

71. Taheri-Talesh, N. et al. The tip growth apparatus of Aspergillus nidulans. Mol Biol Cell 19, 1439–1449 (2008).

72. Delgado, M. et al. The actin module of endocytic internalization in <EM>Aspergillus nidulans</EM>: a critical role of the WISH/DIP/SPIN90 family protein Dip1. bioRxiv, 2025.2002.2013.638025 (2025).

73. Pinar, M. et al. The type V myosin-containing complex HUM is a RAB11 effector powering movement of secretory vesicles. iScience 25, 104514 (2022).

74. Xie, Y. et al. Polarisome scadolder Spa2-mediated macromolecular condensation of Aip5 for actin polymerization. Nat Commun 10, 5078 (2019).

75. Lawson, M.J., Drawert, B., Petzold, L. & Yi, T.M. A positive feedback loop involving the Spa2 SHD domain contributes to focal polarization. PLoS One 17, e0263347 (2022).

76. Zheng, P. et al. Spitzenkorper assembly mechanisms reveal conserved features of fungal and metazoan polarity scaffolds. Nat Commun 11, 2830 (2020).

77. Bähler, J. et al. Heterologous modules for efficient and versatile PCR-based gene targeting in Schizosaccharomyces pombe. Yeast 14, 943–951 (1998).

78. Zhang, X.R., et al. An improved auxin-inducible degron system for fission yeast. G3 12 (2022).

79. Vjestica, A. et al. A toolbox of Stable Integration Vectors (SIV) in the fission yeast Schizosaccharomyces pombe. J Cell Sci (2019).

80. Vjestica, A., Merlini, L., Dudin, O., Bendezu, F.O. & Martin, S.G. Microscopy of Fission Yeast Sexual Lifecycle. J. Vis. Exp. (2016).

81. Thevenaz, P., Ruttimann, U.E. & Unser, M. A pyramid approach to subpixel registration based on intensity. IEEE Trans Image Process 7, 27–41 (1998).

82. Miura, K. Bleach correction ImageJ plugin for compensating the photobleaching of time-lapse sequences. F1000Res 9, 1494 (2020).

83. Sbalzarini, I.F. & Koumoutsakos, P. Feature point tracking and trajectory analysis for video imaging in cell biology. J Struct Biol 151, 182–195 (2005).

